# A basally active cGAS-STING pathway limits SARS-CoV-2 replication in a subset of ACE2 positive airway cell models

**DOI:** 10.1101/2024.01.07.574522

**Authors:** Maritza Puray-Chavez, Jenna E. Eschbach, Ming Xia, Kyle M. LaPak, Qianzi Zhou, Ria Jasuja, Jiehong Pan, Jian Xu, Zixiang Zhou, Shawn Mohammed, Qibo Wang, Dana Q. Lawson, Sanja Djokic, Gaopeng Hou, Siyuan Ding, Steven L. Brody, Michael B. Major, Dennis Goldfarb, Sebla B. Kutluay

**Affiliations:** Department of Molecular Microbiology, Washington University School of Medicine, St. Louis, MO, USA; Department of Cell Biology and Physiology, Washington University School of Medicine, St. Louis, MO, USA; Division of Pulmonary and Critical Care Medicine, Department of Medicine, Washington University School of Medicine, St. Louis, MO, USA; Department of Otolaryngology, Washington University School of Medicine, St. Louis, MO, USA; Institute for Informatics, Data Science & Biostatistics, Washington University School of Medicine, St. Louis, MO, USA

**Author notes:** Correspondence (S.B.K.).

## Abstract

Host factors that define the cellular tropism of SARS-CoV-2 beyond the cognate ACE2 receptor are poorly defined. Here we report that SARS-CoV-2 replication is restricted at a post-entry step in a number of ACE2-positive airway-derived cell lines due to tonic activation of the cGAS-STING pathway mediated by mitochondrial DNA leakage and naturally occurring cGAS and STING variants. Genetic and pharmacological inhibition of the cGAS-STING and type I/III IFN pathways as well as ACE2 overexpression overcome these blocks. SARS-CoV-2 replication in STING knockout cell lines and primary airway cultures induces ISG expression but only in uninfected bystander cells, demonstrating efficient antagonism of the type I/III IFN-pathway in productively infected cells. Pharmacological inhibition of STING in primary airway cells enhances SARS-CoV-2 replication and reduces virus-induced innate immune activation. Together, our study highlights that tonic activation of the cGAS-STING and IFN pathways can impact SARS-CoV-2 cellular tropism in a manner dependent on ACE2 expression levels.

## INTRODUCTION

Severe acute respiratory syndrome coronavirus 2 (SARS-CoV-2) is continuing to cause significant respiratory disease and mortality in vulnerable populations. The emergence of highly transmissible viral variants resistant to vaccines and existing immunity is an ongoing concern. The innate immune response mediated by type I, type II and type III interferons (IFNs) constitutes a potent block to SARS-CoV-2 replication ^1–3^. It remains unknown if IFN-mediated antiviral responses can modulate the cellular tropism of SARS-CoV-2 independently of the cognate host cell SARS-CoV-2 receptor, angiotensin converting enzyme 2 (ACE2). Given the multiple ways in which SARS-CoV-2 can antagonize IFN production and signaling ^1–3^, a comprehensive understanding of the interplay between these mechanisms may inform the observed multi-organ involvement, reveal new facets of emerging SARS-CoV-2 variants and ultimately identify new opportunities for therapeutic intervention.

SARS-CoV-2 homotrimeric spike (S) glycoprotein binding to the host cell ACE2 receptor mediates viral entry ^4–7^. Endogenous ACE2 expression is surprisingly low in the respiratory tract ^8,9^ compared with higher levels in other tissues of the gastrointestinal tract, kidney and myocardium ^10,11^. Although ectopic expression of ACE2 is sufficient for viral entry in several cell culture and animal models ^4–7,12^, additional factors and host cell proteases including TMPRSS2 may contribute to defining the cellular tropism of SARS-CoV-2 (and variants thereof) at physiological levels of ACE2 expression ^13^. Growing evidence suggests that viral attachment factors can facilitate ACE2-dependent and even enable ACE2-independent entry ^13,14^. Together, these alternative receptors and attachment factors may compensate for the low levels of ACE2 expression in various organs, including the distal respiratory system.

In addition to the identity and expression of entry factors and receptors, SARS-CoV-2 cellular tropism can in principle be determined by the basal activation of cell intrinsic and innate immune defenses that are canonically mounted in response to pathogens. For example, a preactivated antiviral innate immunity in the upper airways of young children has been proposed to limit SARS-CoV-2 replication ^15^. In addition, healthy microbiota and endogenous retrovirus activation can also promote antiviral innate immunity through cGAS/STING and subsequent type I/III IFN pathway activation ^16–21^. Interestingly, disruptions in microbiota homeostasis in both the respiratory and the gastrointestinal tracts, the primary targets of SARS-CoV-2 ^22–25^, have been observed in hospitalized COVID-19 patients ^26–30^ highlighting a possible link between basal innate immune activation and SARS-CoV-2 pathogenesis.

Given the potent antiviral effects of type I/III IFNs, numerous SARS-CoV-2 encoded proteins have evolved to antagonize IFN synthesis or subsequent IFN signaling pathways to allow efficient viral spread ^1–3^. On the other hand, we and others have shown that the RIG-I-like receptors (RLRs), RIG-I and MDA5, can detect SARS-CoV-2 replication intermediates ^31–37^, resulting in robust, albeit possibly delayed IFN-dependent innate immune activation ^38–42^. Of note, severe COVID-19 patients also exhibit an impaired type I and III IFN response but a heightened inflammatory response ^43,44^ in part due to the presence of autoantibodies against type I IFNs ^45^. Similarly, young infants exhibit restrained IFN-γ production following SARS-CoV-2 infection compared to adults ^46^. Given these multifactorial antagonistic relationships between SARS-CoV-2 and host cells, cell and tissue tropism and subsequent pathogenesis of SARS-CoV-2 is plausibly shaped by a complex interplay of receptor presence and expression levels, innate immune activation state of the target cells and distinct viral proteins that antagonize the host cell antiviral responses.

In this work, we sought to understand the molecular basis of an intrinsic resistance to SARS-CoV-2 replication in a panel of lung and upper airway cell lines expressing comparably high levels of endogenous ACE2/TMPRSS2^31^ (from herein referred to as ACE2(high)). Analysis of the transcriptomic signatures of 10 airway-derived cell lines reveal markedly higher levels of ISGs and IFN pathway mediators in the ACE2(high) cells. A targeted genetic screen in a representative ACE2(high) cell line reveals that the constitutive activation of the cGAS-STING pathway underlies the observed IFN pathway activity. Genetic disruption and pharmacological inhibition of the cGAS-STING and IFN pathways substantially decrease baseline ISG expression and enhance virus replication in multiple ACE2(high) cell lines. We demonstrate that the basal cGAS-STING pathway activity in these cells is likely maintained by a combination of mitochondrial DNA (mtDNA) leakage and naturally occuring cGAS/STING variants that are more prone to activation. ACE2 overexpression efficiently overcomes the IFN-mediated blocks to SARS-CoV-2 replication, demonstrating the ability of SARS-CoV-2 to overcome these possibly saturable post-entry blocks to infection. SARS-CoV-2 replication in STING knockout cells and in primary human airway cultures triggers type I IFN pathway activation, but only in uninfected bystander cells, suggesting efficient antagonism of the IFN signaling pathway in infected cells. Furthermore, in primary human airway cultures SARS-CoV-2-infected cells display markedly lower levels of STING expression and pharmacological inhibition of STING enhances infectious virion release while limiting innate immune activation. Collectively our study highlights that basal activation of the cGAS-STING and subsequent IFN pathways contribute to defining the SARS-CoV-2 cellular tropism *in vitro* in a manner dependent on ACE2 expression levels. Our findings also establish a pivotal role for STING in establishment of antiviral responses in the relevant airway infection models and expand the culture models available to study CoV-host interactions.

## RESULTS

### SARS-CoV-2 replication is blocked at a post-entry stage in a subset of airway-derived cell lines that endogenously express ACE2

We previously screened 10 human lung and upper airway cell lines expressing variable levels of endogenous ACE2/TMPRSS2 for their ability to support SARS-CoV-2 replication^31^. Unexpectedly, we found four cell models—SCC25, H596, OE21 and Detroit 562—that express comparably higher levels of ACE2 mRNA (**Fig. S1a**) and protein but were resistant to SARS-CoV-2 replication ^31^. Of note, these initial screens were conducted with a SARS-CoV-2 stock carrying the S E484D substitution, which arises consistently during cell culture adaptation and allows virus replication in an ACE2-independent manner ^31,47^. Given this possible complication, we first validated the ability of virus stocks verified to bear wild type (WT) S to infect these ACE2 expressing cells. SARS-CoV-2 (2019-nCoV/USA-WA1/2020) WT S productively infected PC-9, KYSE-30 and H1299 cells that express lower levels of ACE2 (**Fig. S1a**) in addition to the control Calu-3 and HCC827 cells, as evident from the substantial increase in the amount of cell-associated viral RNA (**Fig. 1a**), viral N antigen (**Fig. S1b**) and infectious virion release in cell culture supernatants (**Fig. 1b**) following virus inoculation. CRISPR-mediated knockout of ACE2 decreased cell-associated SARS-CoV-2 RNA by ∼20 to 4000-fold in these cells (**Fig. 1c**), validating the ACE2-dependence of virus replication. In agreement with our prior findings ^31^, ACE2(high) cell lines including SCC25, H596, OE21 and Detroit 562 were largely resistant to SARS-CoV-2 replication (**Fig. 1a-c, S1b**). Importantly, we found that ACE2 knockout led to a significant reduction in incoming cell-associated SARS-CoV-2 RNA in the ACE2(high) cell lines (**Fig. 1d, e**), suggesting a post-binding and/or post-entry block to virus replication and spread. These results demonstrate that ACE2 expression is not sufficient to allow productive SARS-CoV-2 replication in the ACE2(high) SCC25, H596, OE21, and Detroit 562 cell lines.

**Figure 1.**
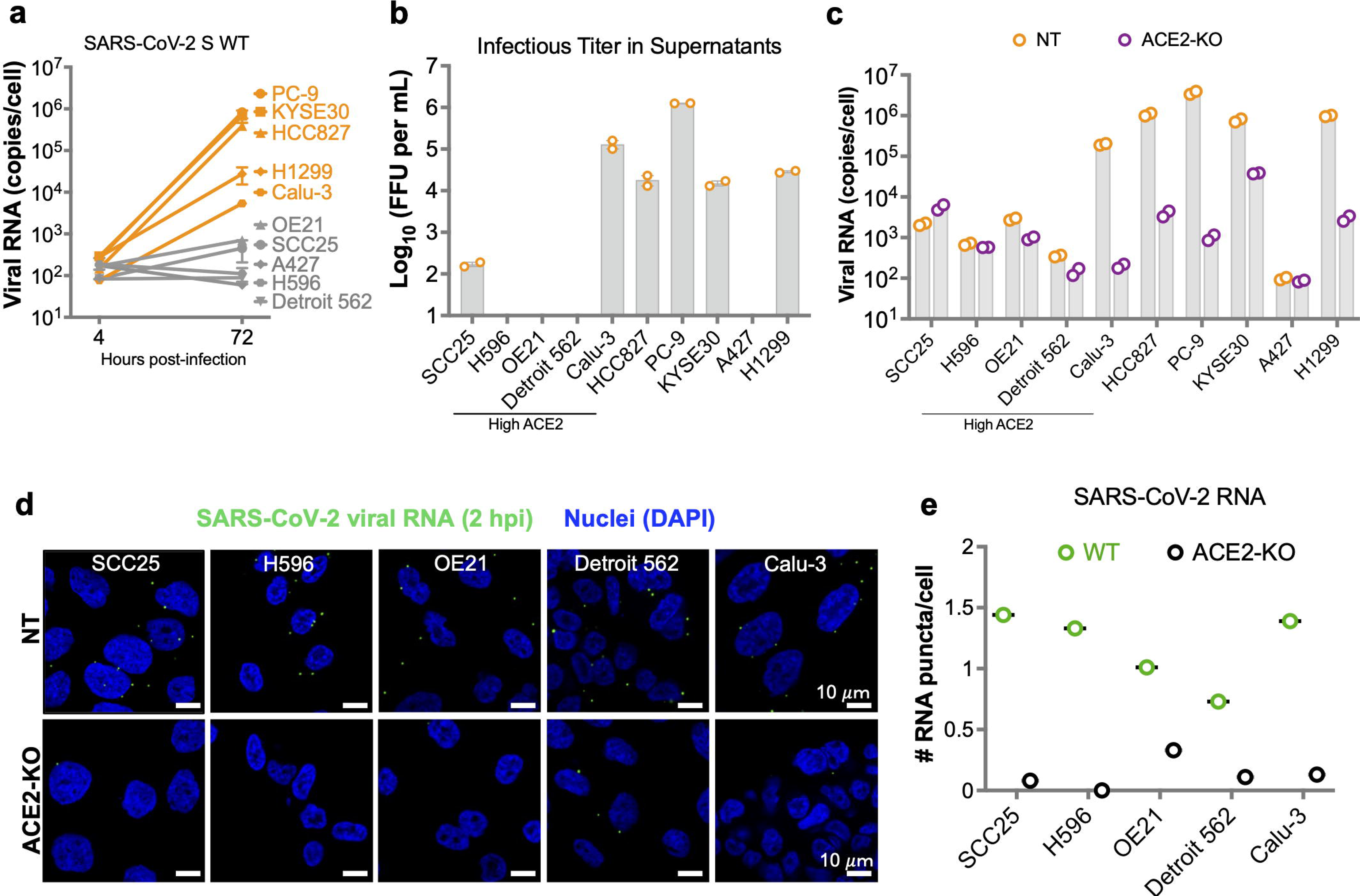
SARS-CoV-2 replication is blocked at a post-entry stage in a subset of airway-derived cell lines that endogenously express ACE2. **(a)** The indicated airway-derived cell lines were inoculated with WT SARS-CoV-2 (2019-nCoV/USA-WA1/2020) at a MOI: 0.15 i.u./cell. RT-qPCR analysis of cell-associated SARS-CoV-2 N sgRNA (copies/cell) at 4 and 72 hpi is shown. Data show the average of 2 independent experiments, errors bars denote the range. **(b)** Focus forming assay performed on culture supernatants of the indicated cells infected with SARS-CoV-2 for 72 hpi (from panel A). Data show focus forming units per mL (FFU/mL, log transformed) from two independent experiments, error bars show the SEM. **(c)** Polyclonal pools of ACE2 knockout (ACE2-KO) and control cell lines transduced with a non-targeting sgRNA (NT) were infected with WT SARS-CoV-2 at a MOI: 2 i.u./cell. RT-qPCR analysis of cell-associated viral RNA (sgRNA N) at 72 hpi is shown from two biological replicates. Data show the mean, error bars show the SEM, **p< 0.01 where significance was assessed by multiple unpaired t-tests. **(d)** Polyclonal pools of ACE2 KO and non-targeting control (NT) cell lines were infected with WT SARS-CoV-2 at a MOI: 2 i.u./cell. In situ hybridization of incoming viral RNA (green) at 2 hpi was performed as detailed in Methods. Nuclear DAPI staining is shown in blue. **(e)** Quantification of data presented in Panel D. Puncta of SARS-CoV-2 RNA was counted in ∼200 cells per cell line from two independent experiments. The average number of RNA puncta per condition is shown in the graph. Source data are provided as a Source Data file.

### ACE2(high) airway-derived cell lines express high baseline levels of IFN pathway genes

A common block to SARS-CoV-2 replication is mediated by type I and type III IFNs ^1–3^, as exemplified by potent inhibition of SARS-CoV-2 replication by IFN-α pretreatment of Calu-3 cells (**Fig. S2a**). Seeking to understand if IFN pathway activity contributed to SARS-CoV-2 resistance in the ACE2(high) SCC25, H596, OE21, and Detroit 562 cells, we first evaluated the expression of IFN pathway response genes by using publicly available RNA-seq data. Strikingly, transcriptomic signatures of the cell line panel revealed substantially higher basal expression levels of numerous ISGs, IFN pathway components and inflammatory mediators (e.g. *CCL2*, *CCL5*, *IL6*, *CXCL9*) in the ACE2(high) cells compared to those that express low to no ACE2 (**Fig. 2a**). RT-qPCR and immunoblotting confirmed this observation and showed markedly high baseline expression of several ISGs and IFN-pathway mediators (i.e. MX1, IFIT1, STAT1, pSTAT1, IRF7 and IRF9) in SCC25, H596 and OE21 cells, moderately higher levels of distinct ISGs in Detroit 562 cells (**Fig. 2b-f, S2b, S2c**). Further investigation of the RNA-seq data revealed elevated levels of type I and type III IFNs in the ACE2(high) SCC25, H596, OE21 but not Detroit 562 cells or those permissive to infection (**Fig. S2d**); however, the amount of IFN-β and IFN-γ protein in cell culture supernatants was below the limit of detection in ELISA assays (**Fig. S2e**, **S2f**). In addition to the basally high ISG levels and despite the lack of noticeable virus replication, SCC25, H596, OE21 and Detroit 562 cells responded to SARS-CoV-2 inoculation by further upregulation of ISGs (**Fig. 2g, 2h, S2g-j**). Collectively, these findings suggest that the basally high levels of ISG expression, coupled with further elevation of ISGs upon SARS-CoV-2 inoculation, may underlie the post-entry block to infection in the ACE2(high) SCC25, H596, OE21 and Detroit 562 cells.

**Figure 2.**
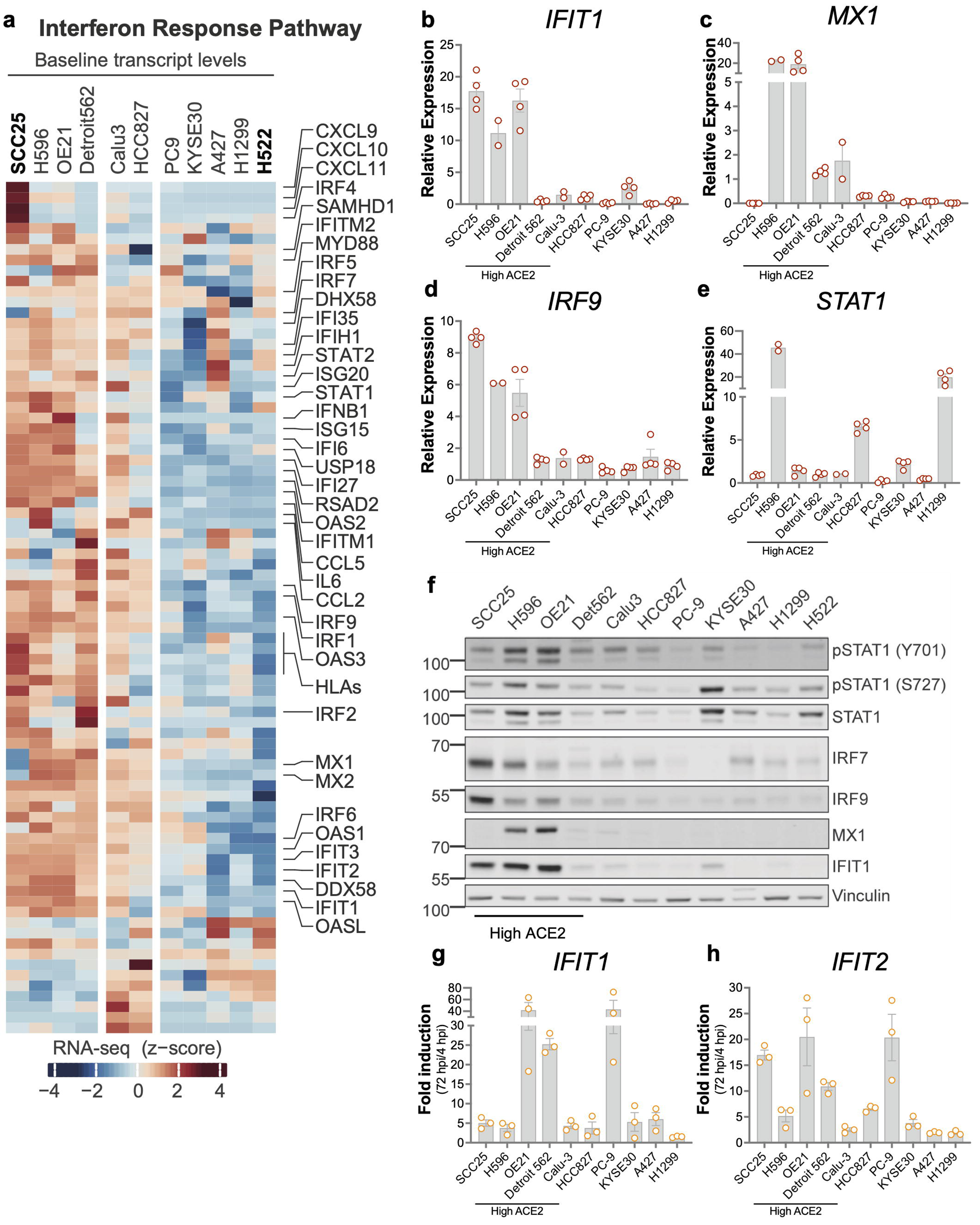
ACE2(high) airway-derived cell lines that are resistant to SARS-CoV-2 replication express high baseline levels of type I IFN pathway genes. **(a)** Hierarchical clustering of IFN response pathway RNA expression levels downloaded from CCLE. The heatmap displays z-scored log_2_(TPM) values. **(b-e)** RT-qPCR analysis of representative ISGs including *IFIT1, MX1, IRF9* and *STAT1* in uninfected cells. Expression levels are normalized relative to Calu-3 cells (set to 1). Data show the mean from n=2 (H596 and Calu-3) or 4 (other cells) independent biological replicates, error bars denote SEM. **(f)** Immunoblot showing pSTAT1 (Y701 and S727), STAT1, IRF7, IRF9, MX1, IFIT1 and Vinculin levels across airway-derived cell lines (not infected). Blots are representative of two independent experiments. **(g, h)** Airway cell lines were infected with SARS-CoV-2 at a MOI of 2 i.u./cell and expression of *IFIT1* and *IFIT2* in was analyzed by RT-qPCR at 4 and 72 hpi. Y-axes indicate ISG fold induction at 72 hpi relative to 4hpi. Data show the mean from n=3 independent replicates, errors bars show the SEM. Source data are provided as a Source Data file.

### Basally active type I/III IFN signaling pathway underlies the resistance of ACE2(high) cell lines to SARS-CoV-2 replication

To evaluate whether the constitutive activity of the canonical type I/III IFN pathway underlies the resistance of SCC25, H596, OE21 and Detroit 562 cells to SARS-CoV-2 infection, cells were pretreated with ruxolitinib, a JAK1/2 inhibitor, and inoculated with SARS-CoV-2 in its presence. Ruxolitinib pretreatment decreased the baseline expression of five canonical ISGs (*IFIT1*, *MX1, IFIT2, ISG15* and *IRF9*) in SCC25 cells (**Fig. 3a, 3b, S3a-c**), though the effects were relatively modest for certain ISGs. Subsets of these ISGs were also suppressed in the other ACE2(high) cell models (**Fig. 3a, 3b, S3a-c**). Ruxolitinib treatment resulted in an approximate 3-log increase in the amount of cell-associated SARS-CoV-2 RNA in OE21 and SCC25, but not in Detroit 562 or H596 cells (**Fig. 3c**). Because of this, we decided to focus our studies primarily on SCC25 cells with validation experiments in OE21 cells. In agreement with the ruxolitinib data, CRISPR-mediated knockout of IFNAR and STAT1 (**Fig. S3d**), and siRNA-mediated depletion of downstream IFN pathway components STAT2 and IRF9 (**Fig. S3e, S3f**) enhanced SARS-CoV-2 replication in SCC25 cells (**Fig. 3d-G, S3g**). STAT1 knockout also increased cell-associated viral RNA levels in OE21 and more modestly in Detroit 562 cells, whereas H596 cells remained resistant to infection despite a reduction in ISG expression (**Fig. S3h-k**). Enhancement of SARS-CoV-2 replication in SCC25 and OE21 cells correlated with a reduction in baseline ISG expression (**Fig. S3l, S3m**), though we noted that expression of certain ISGs including *IFIT2* and *OASL* was less sensitive to STAT1 depletion. Together, these data suggest that the high baseline activity of the canonical type I/III IFN signaling underlies the inability of OE21, SCC25 and Detroit 562 cells to support SARS-CoV-2 replication.

**Figure 3.**
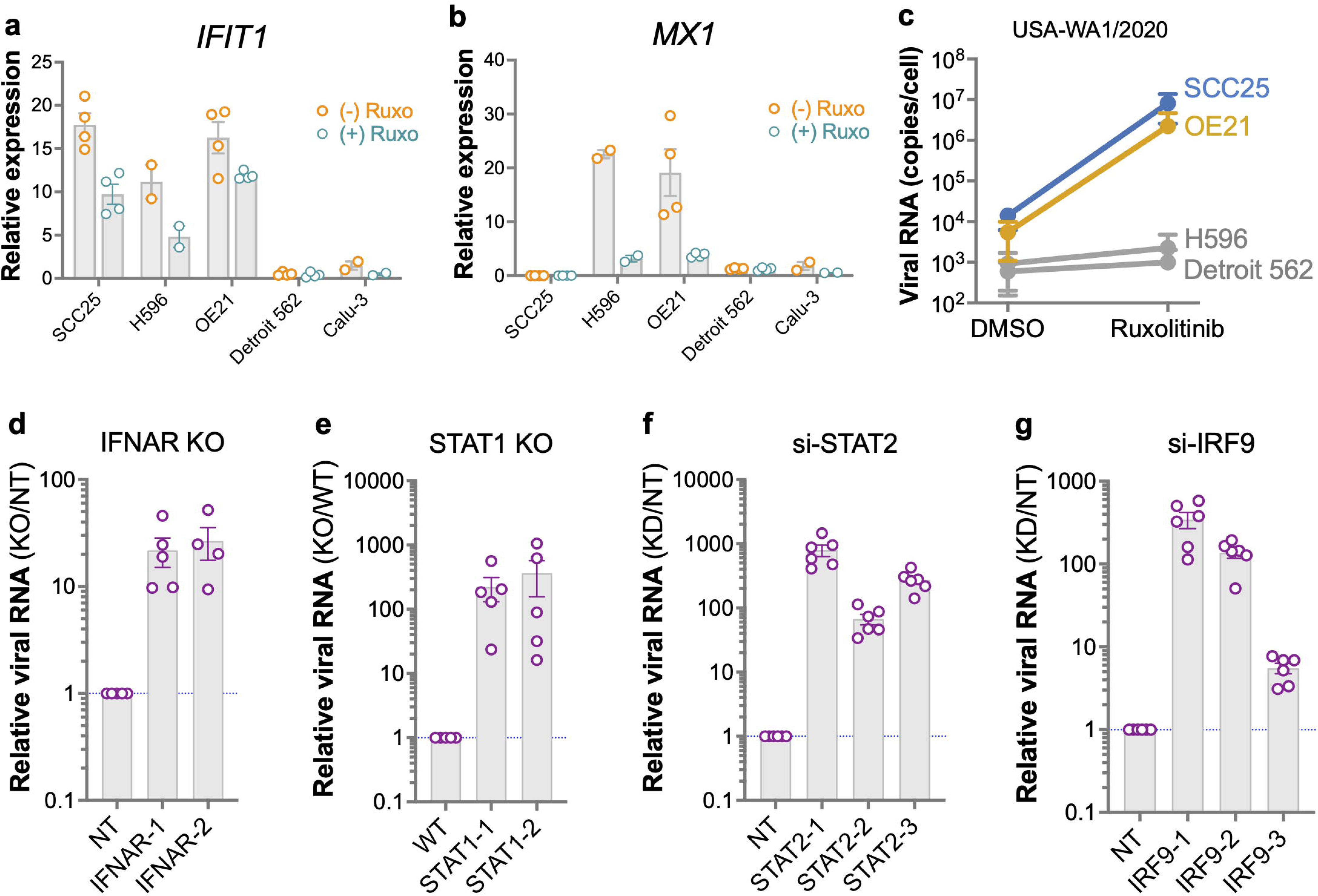
Basally active type I IFN pathway limits SARS-CoV-2 replication in a subset of ACE2(high) cells. **(a, b)** SCC25, H596, OE21, Detroit 562 and Calu-3 cells were pretreated with 1 μM ruxolitinib for 24 hours. *IFIT1* (**a**) and *MX1* (**b**) expression was analyzed by RT-qPCR and expression levels are shown relative to untreated Calu-3 cells (set to 1). Data show the mean from n=2 (H596 and Calu-3) and 4 (other cells) independent biological replicates, error bars show the SEM. **(c)** SCC25, H596, OE21 and Detroit 562 cells were pretreated with DMSO (mock) or 1 μM ruxolitinib for 24 hours and infected with SARS-CoV-2 at a MOI of 2 i.u./cell in presence of DMSO or 1 μM ruxolitinib. RT-qPCR analysis shows cell-associated viral RNA levels at 72 hpi. Data show the mean from n=4 independent replicates, error bars show the SEM. **(d-g)** Polyclonal populations of SCC25 cells knocked out (KO) for IFNAR (**d**) or STAT1 (**e**) using two independent sgRNAs or transduced with a non-targeting (NT) control and SCC25 cells transfected with 3 separate siRNAs targeting STAT2 (**f**), IRF9 (**g**), or a non-targeting control (NT) were infected with SARS-CoV-2 at a MOI of 2 i.u./cell. Cell-associated viral RNA was analyzed by RT-qPCR at 72 hpi. Y-axes indicate the relative expression ratio between knockdown (KD) or knockout (KO) and the respective NT samples. Data display the mean of n=5 (**d, e**) and n=6 (**f, g**) independent replicates, error bars denote the SEM. Source data are provided as a Source Data file.

### Tonic activation of the cGAS-STING pathway underlies the resistance of SCC25, OE21 and Detroit 562 cells to SARS-CoV-2 replication

To identify the molecular basis of the basally high IFN pathway activity in the SCC25 cells, we conducted an siRNA screen (2 siRNAs/target) targeting host cell factors involved in sensing of virus replication intermediates including TLRs (e.g. TLR3, TLR7, TLR8 and TLR9), RLRs (e.g. RIG-I, MDA5 and LGP2) as well as their downstream adaptor molecules (e.g. MyD88, TRIF and MAVS), NOD-like receptor family members (i.e. NOD2, NLRP1, NLRP3 and NLRC5) and the cytosolic DNA sensor STING. Remarkably, depletion of STING by two independent siRNAs substantially increased (∼2-3-logs) cell-associated viral RNA levels in SCC25 cells (**Fig. 4a**). Although knockdown of other sensors and downstream adaptors by individual siRNAs increased viral RNA levels up to 10-fold, effects were comparably modest and siRNA-specific (**Fig. 4a**). Consistent with the screening results, CRISPR-mediated knockout of cGAS, STING as well as downstream signaling molecules TBK1 and IRF3 significantly increased cell-associated viral RNA and antigen levels in SCC25 (**Fig. 4b, 4c**) as well as OE21 cells (**Fig. S4a**). Constitutive cGAS-STING pathway activity was also evident in the higher levels of phosphorylated STING and IRF3 present in SCC25 and OE21 cell lysates (**Fig. S4b**). STING knockout also enhanced SARS-CoV-2 replication in the ACE2(high) Detroit 562 cells, whereas H596 cells remained resistant to infection (**Fig. S4c, S4d**). Furthermore, STING knockout did not enhance virus replication further in the otherwise permissive cells (**Fig. S4c, S4d**).

**Figure 4.**
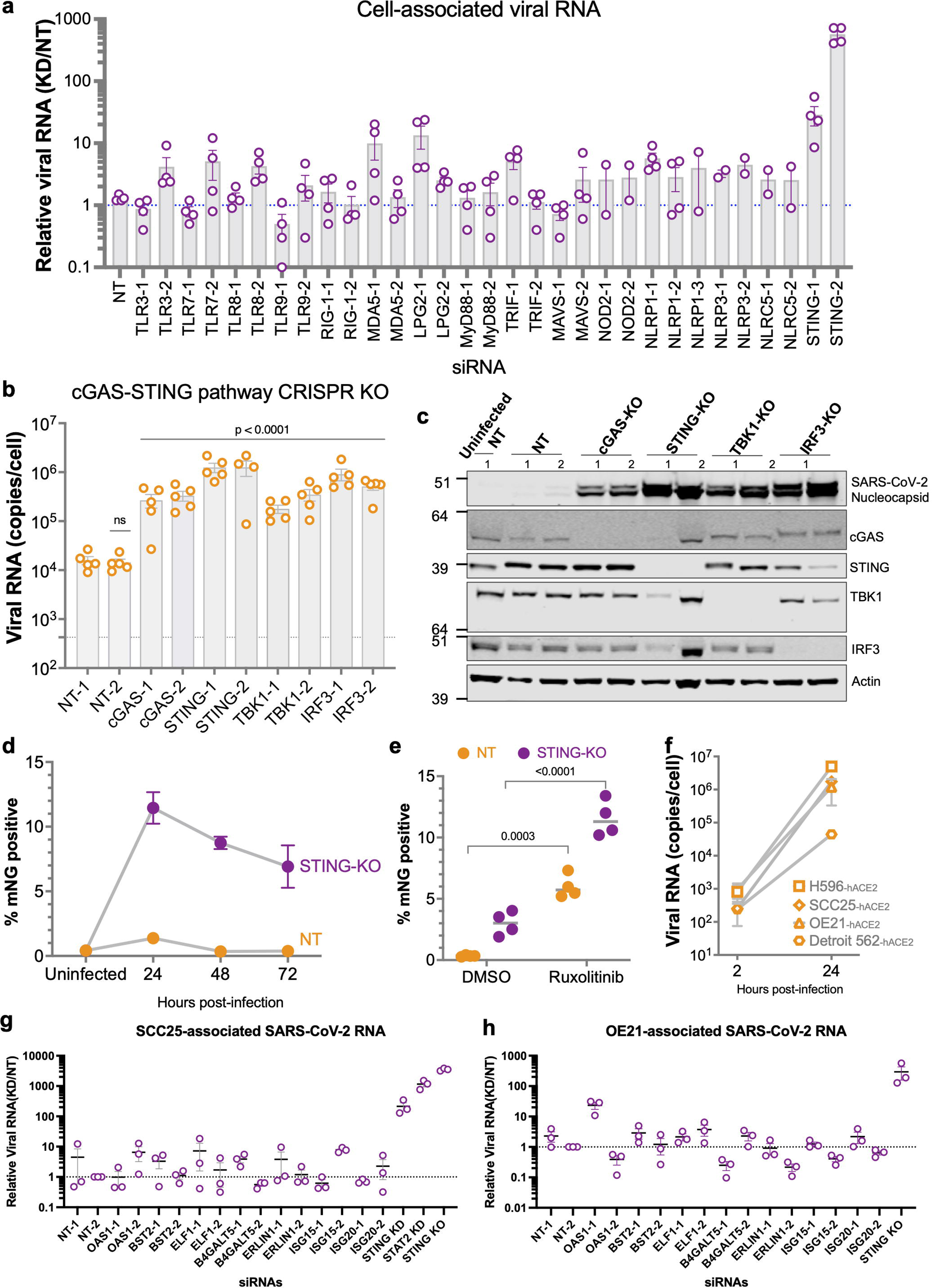
A constitutively active cGAS-STING pathway limits SARS-CoV-2 replication in SCC25 cells. **(a)** SCC25 cells transfected with siRNAs or a non-targeting control (NT) were infected with SARS-CoV-2 at an MOI of 2 i.u./cell for 72 h. Data show the ratio of cell-associated viral RNA between knockdown (KD) and NT samples. Data display the mean of 2 (NLRP1-3, NLRP3-1, NLRP3-2, NLRC5-1, NLRC5-2) or 4 (other targets) biological replicates, with error bars representing SEM. **(b)** SCC25 cGAS, STING, TBK1 and IRF3 KO cells or NT controls were infected with SARS-CoV-2 at MOI: 2 i.u./cell for 72h. Cell-associated viral N RNA from 5 replicates is shown, error bars show the SEM. *p* < 0.0001 significance (compared to NT-1) was assessed by a one-way ANOVA with Dunnett correction. **(c)** Immunoblots (representative of n=2) show SARS-CoV-2 N, cGAS, STING, TBK1, IRF3 and β-actin expression levels in SCC25 cells from panel B. **(d)** SCC25 STING KO cells were infected with SARS-CoV-2-mNG at a MOI:2 i.u./cell. NG-positive cells were enumerated by flow cytometry at 24, 48 and 72 hpi and represented as percentage of total population. Data show the mean of n=5 independent experiments, error bars denote the SEM. **(e)** SCC25 STING KO and NT control cells were pretreated for 24 hours and infected (MOI:2 i.u./cell) in the absence (DMSO) and presence of 1 μM ruxolitinib with SARS-CoV-2-mNG. NG-positive cells were enumerated by flow cytometry at 72 hpi and represented as percentage of total population. Data show the mean of n=4 independent experiments, error bars denote the SEM. *P* values are assessed by two-way ANOVA and Šidák multiple test correction. (**f**) Cells ectopically expressing ACE2 were infected at a MOI: 0.1 i.u./cell with SARS-CoV-2 WT. Cell-associated SARS-CoV-2 RNA at 2 and 24 hpi is shown from n=2 replicates. Data show the mean, error bars denote SEM. (**g, h**) SCC25 (**g**) and OE21 (**h**) cells were transfected with 2 distinct siRNAs or a NT control, infected with SARS-CoV-2 and analyzed as in panel A. Data display the mean of n=3 biological replicates, with error bars representing SEM. Source data are provided as a Source Data file.

Despite the enhanced SARS-CoV-2 replication in STING KO SCC25 cells, we noted that virus spread was limited at later time points in infection (**Fig. 4d, S4e**). Ruxolitinib treatment of STING KO cells allowed greater virus spread (**Fig. 4e**), suggesting that SARS-CoV-2-induced innate immune activation independent of STING limits viral spread. Interestingly, we found that the repressive effects of the type I/III IFN pathway were overcome by ACE2 overexpression in SCC25, OE21, H596 and Detroit 562 cells (**Fig. 4f**) and ruxolitinib treatment did not further enhance virus replication (**Fig. S4f**). It has been reported that the VOCs are generally less sensitive to the inhibitory effects of type I IFNs ^48^. Ruxolitinib pretreatment enhanced virus replication 2-3 logs in SCC25 cells for most of the VOCs with the exception of the Delta and Kappa variant strains, which responded to ruxolitinib treatment only modestly, and the Omicron strain which remained nonresponsive (**Fig. S4g**). We further evaluated whether the IFN-mediated restriction of SARS-CoV-2 replication in SCC25 and OE21 cells was mediated by ISGs known to restrict SARS-CoV-2 replication ^49–51^ through a targeted siRNA screen. While knockdown of certain ISGs (i.e. OAS1) enhanced cell-associated SARS-CoV-2 RNA levels modestly, these effects were relatively modest compared to STING KD/KO and siRNA-specific for both cell models (**Fig. 4g, h**). Collectively, these results suggest that the basally active cGAS-STING and downstream type I/III IFN signaling pathways underlie the resistance to SARS-CoV-2 infection in several ACE2(high) cell line models through the actions of one or more ISGs. While ablation of this pathway enhances SARS-CoV-2 replication early, induction of the type I/III IFN responses at later time points post-infection limits spreading infection.

### cGAS-STING pathway activity underlies the basally high ISG levels in the ACE2(high) cell models

To directly link the tonic cGAS-STING pathway activity with high basal levels of ISG expression in the ACE2(high) cell models, we first interrogated the gene expression profile of WT and STING KO SCC25, H596, OE21 and Detroit 562 cells by RNA-seq. Of the total of 1740 differentially expressed genes in WT vs. KO conditions in a given cell line (**Supplementary data 1**), genes involved in the IFN response pathways stood out as the most significantly downregulated in STING KO cell lines (**Fig. S5a, S5b, Supplementary data 1**). In total, 71 genes involved in IFN and inflammatory response pathways were significantly downregulated in one or more cell lines upon STING KO (**Fig. 5a, Supplementary data 1**). RT-qPCR analysis of select ISGs in cGAS and STING KO SCC25 cells confirmed these findings (**Fig. S5c**). In agreement with this genetic evidence, pharmacological inhibition of STING by H-151, which blocks the activation-induced palmitoylation and clustering of STING ^52^, and SN-011, which binds to the cyclic dinucleotide (CDN)-binding pocket of STING and locks it in an open inactive conformation ^53^, reduced ISG expression in a dose-dependent manner in ACE2(high) cell lines, with SCC25 and OE21 cells being the most responsive to treatment (**Fig. 5b-g, S5d-g**). Pre-treatment of SCC25 cells with SN-011 was sufficient to enhance SARS-CoV-2 replication, whereas H-151 had more modest effects likely due to less efficient downregulation of ISG expression (**Fig. 5h, 5i**). Expectedly, STING inhibition in SCC25-STING KO cells didn’t further enhance SARS-CoV-2 replication (**Fig. 5h, 5i**). On the other hand, neither SN-011 nor H-151 enhanced SARS-CoV-2 replication in OE21 cells (**Fig. 5h, 5i**), likely due to insufficient inhibition of ISG expression by pre-treatment alone. Taken together, both genetic and pharmacological evidence presented herein supports the finding that tonic activation of the cGAS-STING pathway underlies the basally high ISG expression in multiple airway-derived cell models.

**Figure 5.**
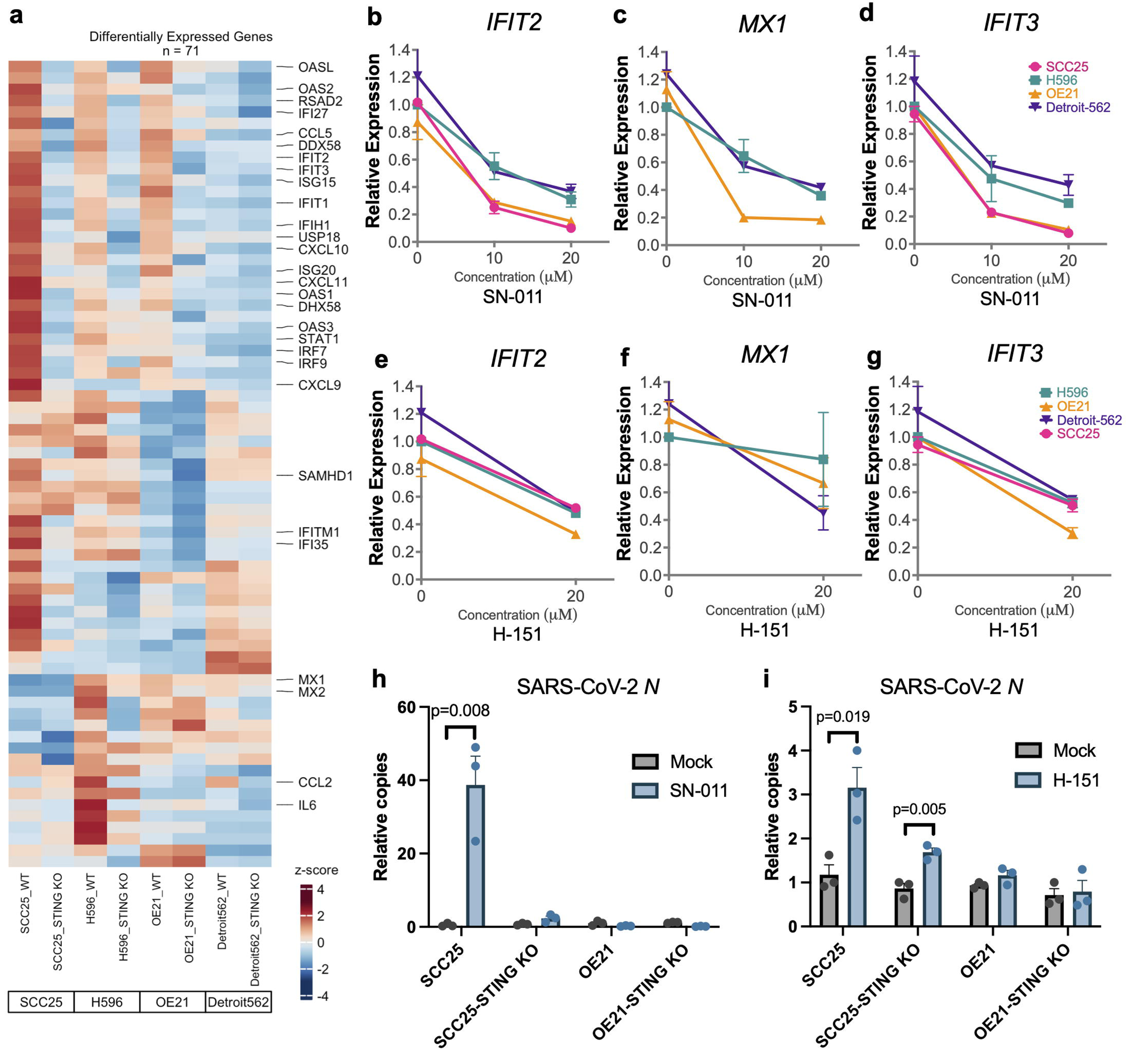
Genetic ablation and chemical inhibition of STING reduces basal ISG levels and enhances SARS-CoV-2 replication. **(a)** WT and STING KO SCC25, H596, OE21 and Detroit 562 cells were subjected to RNA-seq analysis. Heatmap shows differentially expressed genes (absolute log_2_ fold change of >2 and an adjusted *q* value of <0.05 from n=4 independent replicates for each cell type) in the IFN and inflammatory pathways in any given cell line between the WT vs. STING KO variant. (**b-g**) RT-qPCR analysis of *IFIT2, MX1* and *IFIT3* in uninfected SCC25, H596, OE21 and Detroit 562 cells treated with increasing concentrations of SN-011 (**b-d**) and H-151 (**e-g**) for 48 h. Data show the mean from n=2 independent biological replicates, error bars show SEM (**h-i**) WT and STING KO SCC25 and STING KO cells were pre-treated with 20 μM SN-011 (**h**) or H-151 (**i**) for 24h were infected with WT SARS-CoV-2 at MOI: 1 i.u./cell for 72h. Data show the copies of cell-associated SARS-CoV-2 N RNA in compound-treated cells normalized relative to mock-treated samples. Data show the mean from n=3 independent biological replicates, error bars show SEM. Significance was assessed by multiple unpaired two-tailed t-tests and *p* values provided in the figure. Source data are provided as a Source Data file.

### Culture supernatants from and co-culturing with ACE2(high) cell lines induce innate immune activation

Despite the genetic and pharmacological evidence which strongly suggested the involvement of type I/III IFN pathway in mediating resistance of the ACE2(high) cells to SARS-CoV-2 replication, we were unable to detect IFN-β and IFN-γ in culture supernatants. This is possibly due to low level of cGAS-STING pathway activation (as also evident in P-STING and P-IRF3 blots in **Fig. S4b**) and/or possibly unstable nature of secreted IFNs in culture media. To further evaluate the resistance phenotype, we next tested whether culture supernatants from the ACE2(high) cells can activate the IFN pathway in THP-1 human monocytic cells that are highly responsive to type I/III IFNs. Although culture supernatants from SCC25, H596 and OE21 cells trended to trigger ISG expression, as exemplified by the increase in *MX1*, the magnitude of this response was comparably small to that induced by IFN-α treatment but notably higher than the STING KO variants (**Fig. 6a**). Notably, co-culturing of suspension THP-1 cells with SCC25, H596 and OE21 cells but not their STING KO derivatives more potently induced *MX1* expression (**Fig. 6b**). A similar degree of induction was observed in THP-1-cGAS KO cells (**Fig. S6a**), suggesting the independence of IFN pathway activation on possible PAMPs present in SCC25, H596 and OE21 cells/culture medium.

**Figure 6.**
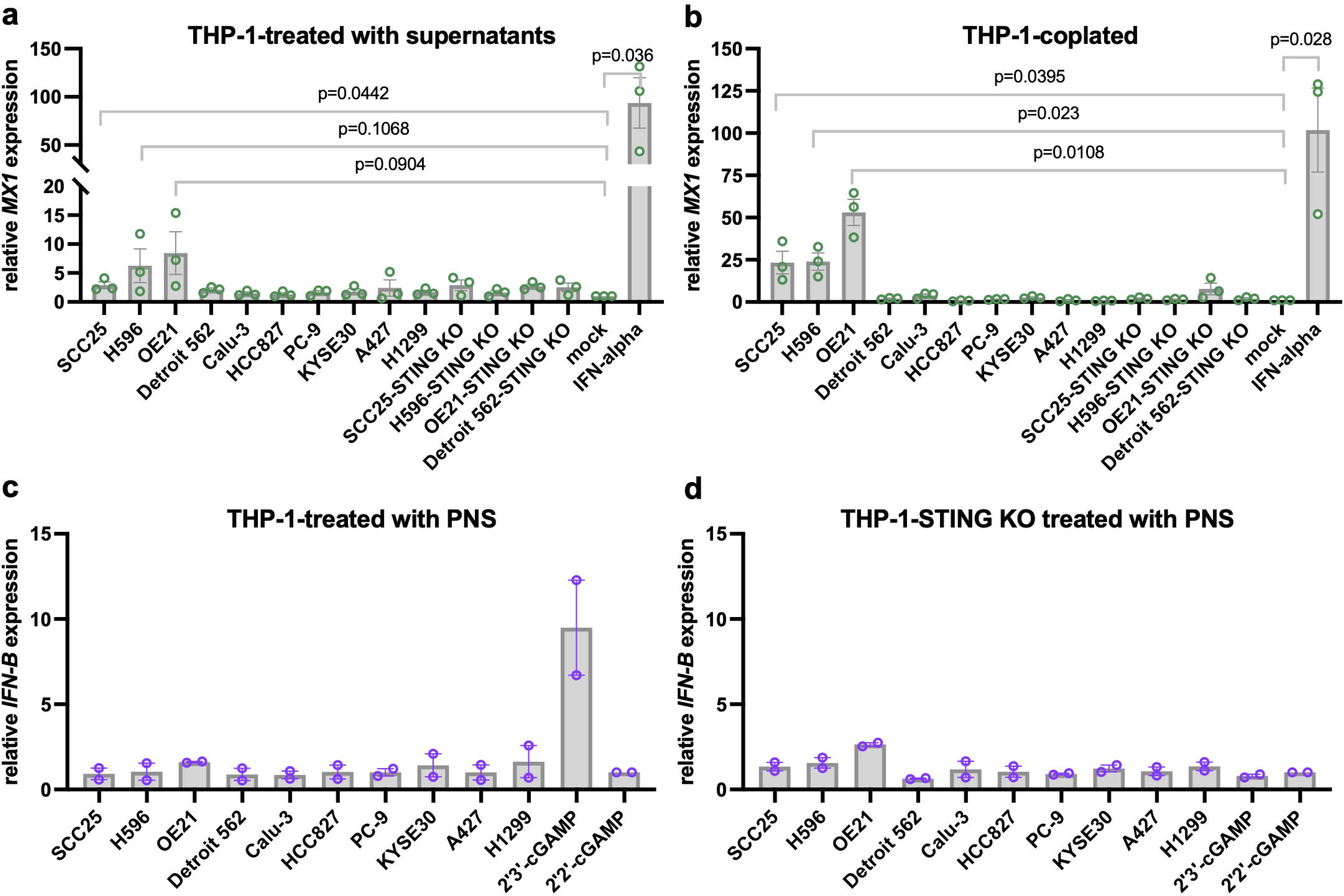
Cell culture supernatants but not cell lysates from SCC25, H596 and OE21 cells trigger innate immune activation. (**a, b**) *MX1* expression was analyzed by RT-qPCR in THP-1 cells treated with cell culture supernatants from (**a**) or co-cultured with (**b**) the indicated cell line panel as explained in Methods. *MX1* expression is normalized relative to mock-treated cells. THP-1 cells treated with 1000 u/ml IFN-α serves as the positive control. Data show the mean induction from n=3 independent biological replicates, error bars display the SEM*. p* values were calculated by multiple unpaired t-tests with Welch’s correction.. (**c, d**) THP-1 or THP-1 STING KO cells were treated with post-nuclear supernatants (PNS) from the indicated cells, 10 μg/ml 2’3’-cGAMP or 10 μg/ml 2’2’-cGAMP for 24 h, as explained in Methods. *IFN-β* expression was analyzed by RT-qPCR and normalized relative to 2’2’-cGAMP-treated samples. Data show the mean from n=2 independent experiments, error bars show the SEM. Source data are provided as a Source Data file.

To evaluate the levels of 2’3’-cGAMP in ACE2(high) cells, we next determined whether post-nuclear fractions from our cell line panel can trigger IFN-β expression in target cells. While, 2’3’-cGAMP, but not 2’2’-cGAMP, induced IFN-β expression over 10-fold in THP-1 and THP-1 cGAS KO cells, PNS from the ACE2(high) cells didn’t (**Fig. 6c, S6b**). Predictably, 2’3’-cGAMP didn’t induce IFN-β expression in the THP-1 STING KO cells (**Fig. 6d**). Overall, these experiments demonstrated that the SCC25, H596 and OE21 cells likely maintain low levels of cGAS-STING-dependent innate immune activation marked with phenotypically detectable levels of type I/III IFN secretion.

### SARS-CoV-2 antagonizes the IFN but not inflammatory pathway activity in infected cells

The limited virus spread in STING KO SCC25 cells (**Fig. 4d, 4e, S4e**) prompted us to further investigate the interplay between cGAS-STING-mediated activation of the IFN pathway and SARS-CoV-2 infection. Despite the noted reduction in ISG expression (**Fig. 5a, S5c, S7a**), cGAS and STING KO cells still mounted substantial increases in ISGs expression in response to SARS-CoV-2 replication (**Fig. 7a**), nearing levels observed in WT SCC25 cells (**Fig. 7b**). In agreement with prior studies ^31–37^, this response was mediated by sensing of SARS-CoV-2 replication intermediates by RIG-I-like receptors, RIG-I and MDA5, whose knockdown abolished ISG induction (**Fig. S7b**). Most notably and in agreement with prior work ^54^, in SARS-CoV-2-infected SCC25-STING KO cells, ISG induction, as exemplified by IFIT3, was largely limited to uninfected bystander cells; cells infected with SARS-CoV-2 were devoid of IFIT3 (**Fig. 7c, S7c, S7d**). Further characterization of single-cell sorted infected and bystander cells revealed a general suppression of ISG expression but elevated expression of numerous inflammatory mediators, including *CXCL1*, *CXCL2*, *CXCL10* and *CCL20* in infected cells (**Fig. 7d, e**). These findings collectively suggest that SARS-CoV-2 can efficiently antagonize IFN signaling but not IFN production or inflammatory pathways in infected cells and that IFN signaling in bystander cells limits viral spread in settings where ACE2 is expressed at endogenous levels.

**Figure 7.**
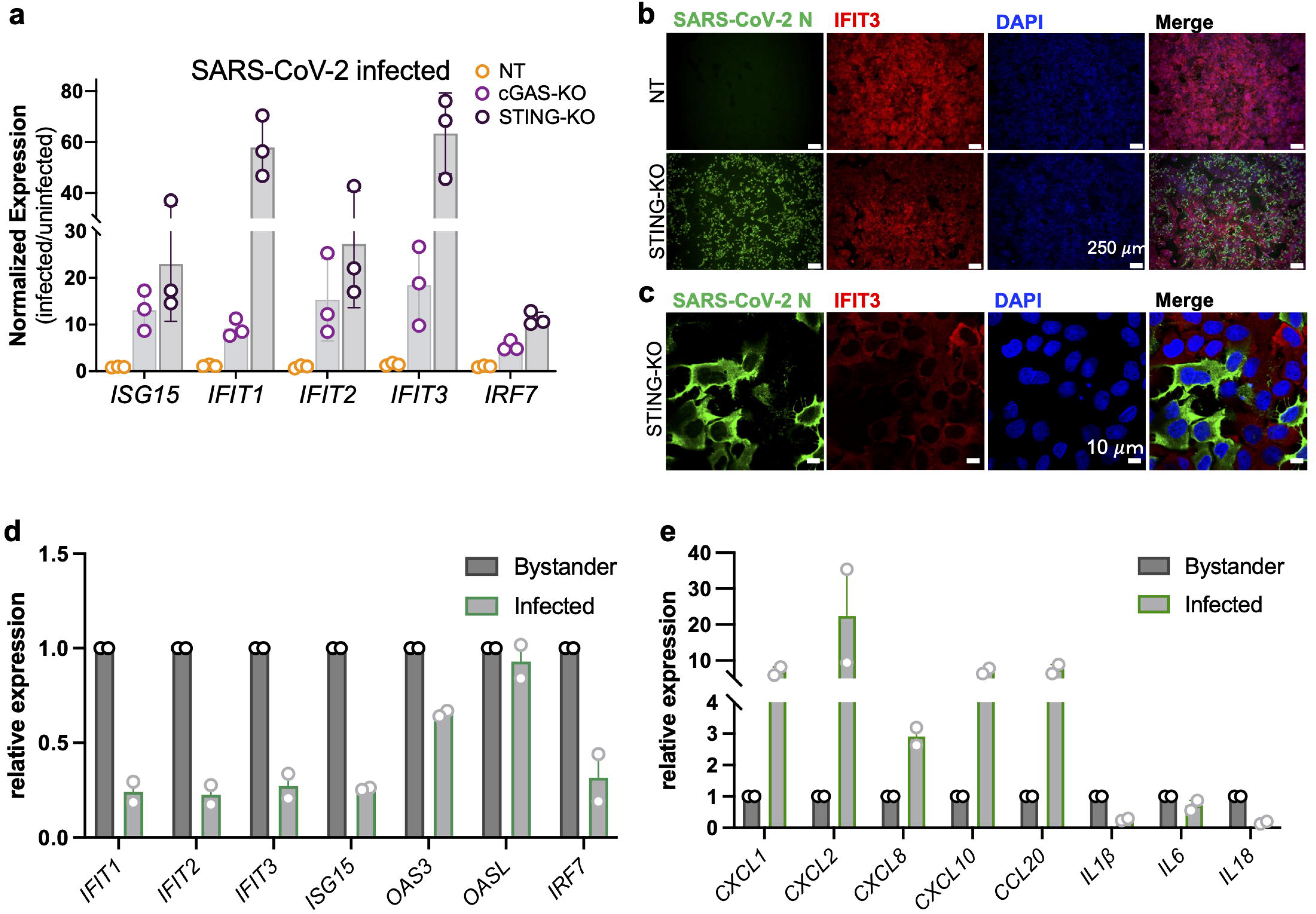
SARS-CoV-2 antagonizes the IFN pathway but not IFN synthesis or inflammatory pathways in infected cells. **(a)** Relative expression of ISGs including *ISG15, IFIT1, IFIT2, IFIT3 and IRF7* in SCC25 cGAS KO, STING KO or NT control cells infected with SARS-CoV-2 at a MOI of 2 i.u./cell. Cells were collected at 72 hpi and ISG expression analyzed by RT-qPCR. Data show the fold induction of ISGs in infected over uninfected cells from n=3 biological replicates, errors bars show the SEM. **(b, c**) SCC25 STING KO or NT control cells were infected with SARS-CoV-2 at an MOI of 2 i.u./cell. Immunofluorescence detection for IFIT3 expression (in red) and SARS-CoV-2 nucleocapsid (N) viral protein (in green), and cellular nuclei (DAPI, in blue) at 72hpi from a representative experiment (n = 2) are shown. Images were collected with an epifluorescence microscope, 4X objective (**b**) or Zeiss LSM 880 Airyscan confocal microscope equipped with a ×63/1.4 objective (**c**) as detailed in Methods. Scale bars = 250 μm (**b**) or 10 μm (**c**). (**d, e**) SCC25 STING KO cells were infected with SARS-CoV-2-mNG at MOI:1 i.u./cell and cells were single-cell sorted at 72 hpi to isolate infected (mNG positive) and bystander cells as detailed in Methods. Expression of the indicated ISGs (**d**) and inflammatory genes (**e**) were analyzed by RT-qPCR in these sorted cell populations. Data show the mean from two independent replicates, error bars show the SEM. Source data are provided as a Source Data file.

### Dissection of mechanisms that maintain tonic cGAS-STING pathway activity

To identify the mechanisms that may underlie tonic activation of the cGAS-STING pathway, we primarily focused on the role of TREX1 endonuclease ^55,56^, possible mutations within components of the cGAS and STING pathway, mitochondrial DNA leakage ^57,58^ and the levels and activities of endogenous retroviruses and transposable elements ^20^. SCC25, H596, OE21 and Detroit 562 cells expressed equivalent levels of TREX1 (**Fig. S8A**) and while TREX1 KO modestly elevated ISG expression in SCC25, H596 and Detroit 562 cells, OE21 TREX1 KO cells displayed surprisingly lower levels of ISGs (**Fig. 8A, S8B**). Interrogation of the RNA-seq data (**Fig. 5A**) revealed that all four cell lines expressed the common H232R variant of STING (**Fig. 8B**), which more potently activates innate immunity than the less common H232 variant ^59,60^. In addition, H596 and Detroit 562 cells expressed the cGAS T35N and T35N/P261H variants, respectively. Of note, cGAS P261H variant has recently been reported to maintain baseline innate immune activation ^61^. Although these naturally occurring variants can contribute to the basal levels of ISGs, we did not identify canonical STING activating mutations such R284M/G or V155M more commonly associated with inflammatory conditions ^62^. No other mutations were identified in TREX1 or ENPP1, the main 2’3’-cGAMP hydrolase. While we did not detect higher levels of cytosolic mtDNA in the ACE2(high) cell lines (**Fig. S8C**), VBIT-4, a VDAC inhibitor that prevents its oligomerization and subsequent mtDNA leakage into the cytosol, diminished ISG expression in SCC25, OE21 and more modestly in Detroit 562 cells, but had no effect on ISG levels in H596 cells (**Fig. 8C**). Furthermore, extended culturing of ACE2(high) cells in the presence of nucleoside analog azidothymidine (AZT), which has been shown to be active against endogenous virus reverse transcriptases ^63^ did not reduce ISG levels (**Fig. 8D, S8D**). In agreement, we did not find an association between cytosolic ERV/LINE DNA (**Fig. 8E**) or RNA levels (**Fig. S8E**) and basally high ISG expression in the cell line panel. Taken together, these findings suggest that mtDNA leakage combined with naturally occurring cGAS and STING variants may explain the tonic cGAS-STING pathway activity in the ACE2(high) cell lines.

**Figure 8.**
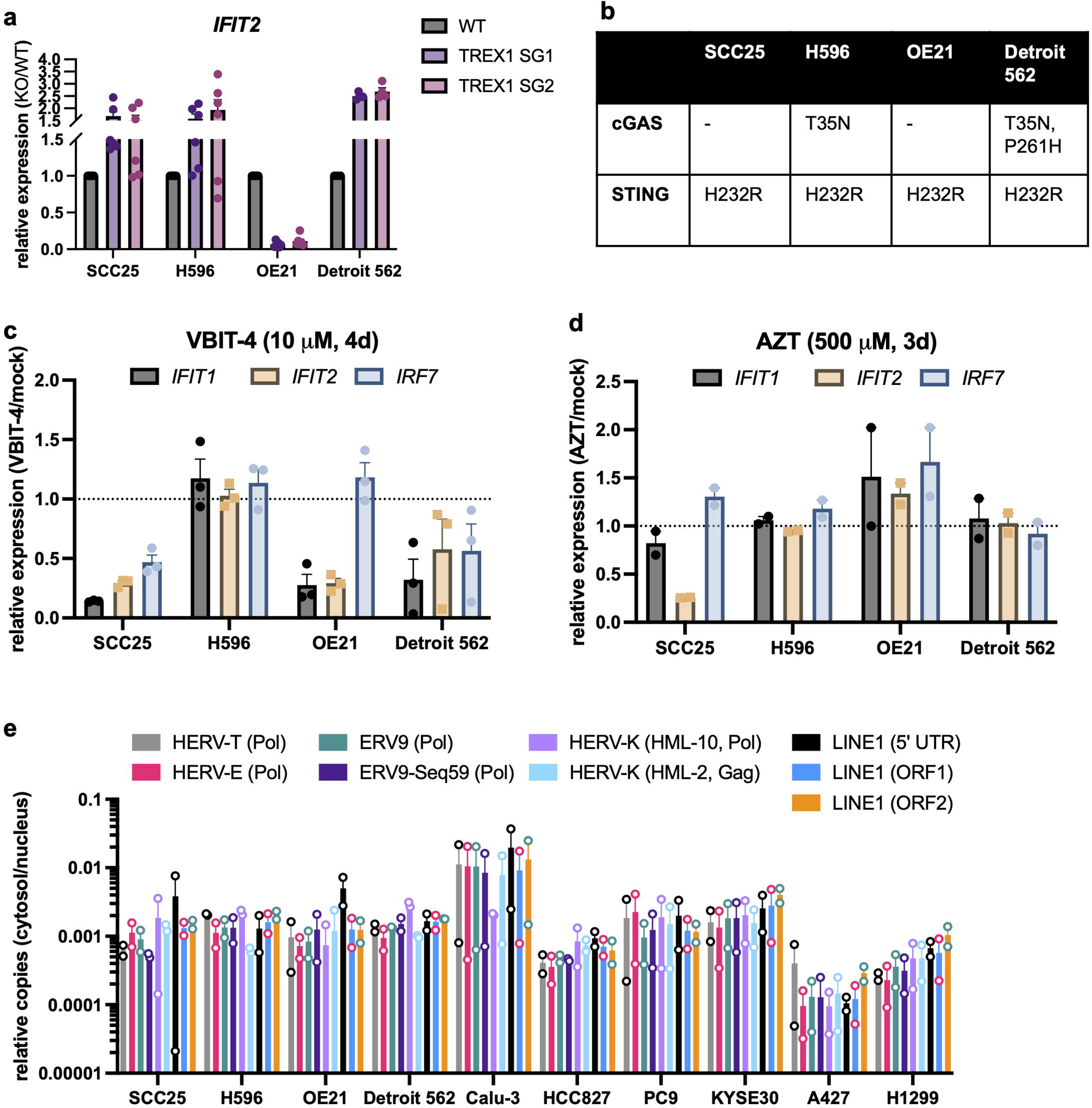
Analysis of mechanisms that underlie tonic cGAS-STING activation in the ISG(high) cell lines. **(a)** Polyclonal populations of SCC25, H596, OE21 and Detroit 562 cells knocked out (KO) for TREX1 using two separate sgRNAs or wild-type (WT) controls were analyzed for *IFIT2* expression by RT-qPCR. Data show the relative expression of *IFIT2* in KO cells compared parental controls (set to 1) from n=3 (OE21) or 6 (SCC25, H596, Detroit 562) replicates, error bars show the SEM. (**b**) cGAS and STING variants present in the indicated cell lines derived from RNA-seq data. (**c, d**) SCC25, H596, OE21 and Detroit 562 cells grown in the presence of 10 μM VBIT-4 for 4 days (**c**) or 500 μM AZT for 3 days (**d**). *IFIT1*, *IFIT2* and *IRF7* expression was analyzed by RT-qPCR and normalized relative to mock-treated controls (set to 1). Data show the mean from n=3 replicates, error bars show the SEM. (**e**) Following subcellular fractionation (see Methods), levels of the indicated HERV and LINE DNA elements were assessed by qPCR in the cytosol and nucleus. Data show the relative levels of each element in the cytosol relative to the nuclear extract in each cell type (n=2, error bars show SEM). Source data are provided as a Source Data file.

### In primary airway cultures, SARS-CoV-2 induces ISG expression in bystander and antagonizes STING expression in infected cells

To extend the findings from our cell line studies to a physiologically relevant model, we next investigated the expression levels of two representative ISGs (i.e. IFIT2, ISG15) and STING in primary human airway epithelial cells (hTECs) grown at the air-liquid interface. Of the five donors analyzed, all except for Donor 2 appeared to express IFIT2 in a subset of cells in the absence of infection (**Fig. 9A**). SARS-CoV-2 infection resulted in an increase in IFIT2 expression in all donors (**Fig.9A**). Remarkably, as in SCC25 STING KO cells, we found that IFIT2 expression was largely limited to uninfected bystander cells (**Fig.9A-C**) and similar results were observed with ISG15 (**Fig. 9D, 9E, S9A**). All five hTECs donors endogenously expressed STING (**Fig. S9B**) and SARS-CoV-2 infection resulted in a modest increase in STING expression (**Fig. 9F, S9C**); however, we found that infected cells were largely devoid of STING (**Fig. 9F, 9G, S9C, S9D**). Inhibition of STING by both SN-011 and H-151 modestly enhanced infectious virion release (**Fig. 9H**), which in the case of H-151 correlated with a decrease in ISG expression in infected hTECs (**Fig. 9I**). In contrast, SN-011 only modestly reduced ISG expression in one Donor and ISG15 expression in all donors (**Fig. S9E**). Analysis of publicly accessible single-cell data sets revealed STING and ISG expression in healthy alveolar type-1 (AT1) human airway cells but not in alveolar type-2 (AT2) cells (**Fig. 10**), which are the primary targets of SARS-CoV-2 ^64^. Together, these findings establish an inhibitory role for STING in SARS-CoV-2 replication and highlight it as a possible factor that contributes to defining the cellular tropism of SARS-CoV-2. These experiments also validate our findings from the cell line models in physiologically relevant settings and demonstrate efficient antagonism of the IFN pathway in productively infected cells.

**Figure 9.**
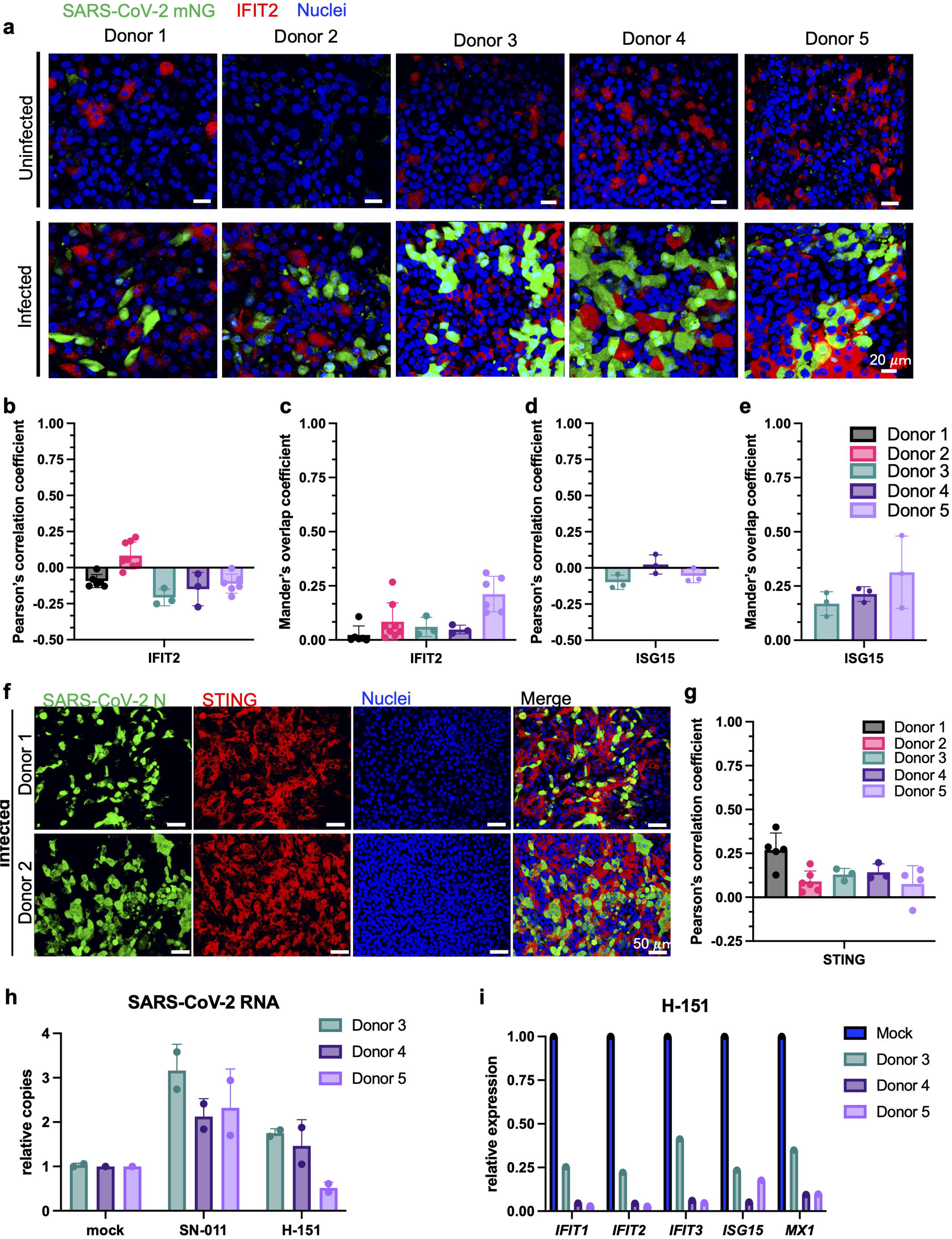
Analysis of ISG and STING expression in SARS-CoV-2-infected primary human airway epithelial cells (hTECs) grown at ALI. **(a-e)** hTECs from five donors grown at ALI were infected with SARS-CoV-2-mNG at a MOI of 2 i.u./cell and fixed at 72 hpi. Immunofluorescence detection for IFIT2 expression (in red), ISG15, mNG (in green), and cellular nuclei (DAPI, in blue) is shown (**a**). Images were analyzed by confocal microscopy with a ×40/1.4 objective. Scale bars = 20 μm, (representative of n=2). Pearson’s correlation (**b, d**) and Mander’s overlap (**c, e**) coefficient derived from imaging experiments. Colocalization analysis of Z stacks was performed using the Colocalization module of Volocity from 3 to 8 separate Z-stacks. Individual numbers plotted display the corresponding values from each Z-stack for each Donor, error bars show the SEM. **(f, g)** hTECs infected as in panel A were stained for STING (in red), SARS-CoV-2 N (in green), and cellular nuclei (DAPI, in blue). Images (**f**) were analyzed by confocal microscopy with a ×20 objective. Scale bars = 50 μm, (representative of n=2). Colocalization analysis of Z stacks was performed as above and Pearson’s correlation coefficient derived from 3 to 6 separate Z-stacks was plotted, error bars show SEM (**g**). (**h, i**) hTECs from three donors were pretreated with 20 μM SN-011 or H-151 for 48h (both apical and basal chambers) and infected with SARS-CoV-2 mNG as above in the presence of compounds in the basal chamber. Virus release in the ALI chamber (**h**) and expression levels of the indicated ISGs (**i**) were analyzed by RT-qPCR. Data in (**h**) show the copies of cell-free SARS-CoV-2 N RNA in culture supernatants of compound-treated cells normalized relative to mock-treated samples. Data show the mean from n=2 independent biological replicates, error bars show SEM. Source data are provided as a Source Data file.

**Figure 10.**
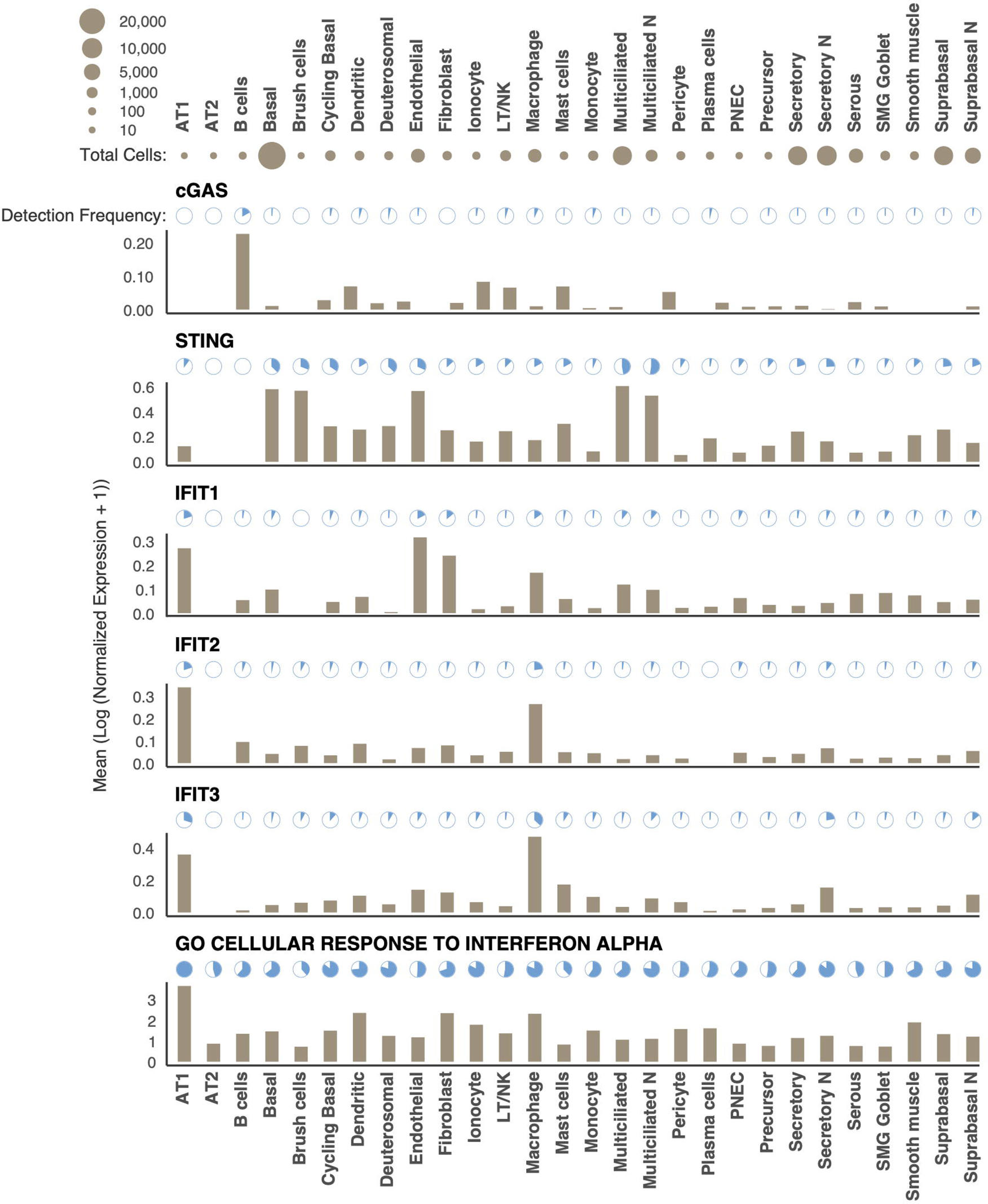
Expression profiles of cGAS, STING and representative ISGs in the healthy human airways. Normalized counts and cell type metadata were downloaded from a publicly available single-cell study ^94^. Detection frequency represents that percentage of cells with > 0 counts for respective gene. Bar plots display the average normalized expression for a gene within a cell type. For the gene set representing the interferon alpha Gene Ontology term, normalized expression levels were summed for all genes in the set prior to computing detection frequencies and average expression.

## DISCUSSION

Because ectopic expression of ACE2 allows efficient SARS-CoV-2 replication in numerous cell lines and animal models ^4–7,12^, it is generally thought that ACE2 is the main factor that defines the cellular tropism of SARS-CoV-2. Our findings challenge this notion and demonstrate that in cells that express endogenous levels of ACE2, baseline activation of the type I/III IFN pathway efficiently limits virus replication at a post-entry step. Interestingly, we found that ectopic expression of ACE2 overcomes the ISG-mediated restriction to SARS-CoV-2 replication in these cell models, suggesting the ability of SARS-CoV-2 to antagonize these blocks at higher multiplicities of infection and possibly their saturable nature. In addition to providing novel insight to molecular mechanisms that govern the cellular tropism of SARS-CoV-2, the ACE2(high) cells studied here can serve as unique models to study the molecular details of innate host responses against SARS-CoV-2.

Our study revealed that the basal activity of cGAS and STING—which have well described roles in sensing of cytosolic DNA including virus replication intermediates ^65,66^—underpins the IFN pathway activation in three separate ACE2(high) cell lines. With regards to what activates the cGAS-STING pathway, we found the involvement of mtDNA leakage in SCC25 and OE21 cells and possibly naturally occurring cGAS and STING mutations that are more prone to activation in the other ACE2(high) cell models. It is, however, important to emphasize that the degree of cGAS-STING pathway activation in the cell lines we studied is likely low and/or present only in a subpopulation of the cells in culture. This would explain why extended treatment with VBIT-4 reduced ISG expression despite the lack of detectable cytosolic mtDNA and 2’3’-cGAMP, the low levels of IFNs present in culture supernatants and phosphorylated STING and IRF3 in cell lysates. What activates the mtDNA leakage is less clear and it is possible that genomic instability commonly present in cancer cell lines ^67^ may also contribute.

Activation of the cGAS-STING pathway has been shown to contribute to SARS-CoV-2 immunopathology and establishment of antiviral defenses ^68^. In macrophages, cGAS activation from engulfed DNA of dying endothelial cells near sites of tissue damage can trigger type I IFN responses leading to lung inflammation in COVID-19 patients ^69^. In addition, in SARS-CoV-2-infected cells both mitochondrial and nuclear DNA leakage into the cytosol can activate cGAS-STING signaling ^69,70^. While STING agonists in uninfected cells can block SARS-CoV-2 infection via triggering type I IFNs ^54,71,72^, pharmacological inhibition of STING reduces severe lung inflammation induced by SARS-CoV-2 and improves disease outcome in mice ^68^. Our study supports the published antiviral functions of cGAS-STING pathway, highlights it as a key factor in mounting the antiviral defenses, and further expands its role to defining the cellular tropism of SARS-CoV-2 *in vitro*. Healthy gut microbiome-induced cGAS-STING activation at a basal state *in vivo* ^19^ represents a particularly relevant context for our observations. SARS-CoV-2 can efficiently infect cells within gastrointestinal tract *in vitro* ^73,74^, but it is possible that microbiota-induced activation of the cGAS-STING pathway can limit SARS-CoV-2 replication within the GI tract or its tropism within particular cell subsets therein. Although STING didn’t appear to be involved in host defense and pathology in a mouse model ^75^, the use of non-native K18 promoter driving ACE2 overexpression in tissues, which we show can decrease sensitivity to IFN-mediated inhibition, is a possible caveat.

Our findings are also suggestive of an antagonistic relationship between SARS-CoV-2 and STING in the primary HBEC-ALI cells; a significant number of productively infected cells lack STING expression (**Fig. 9f, 9g, S9c, S9d**) and STING inhibitors modestly enhanced virus replication while decreasing innate immune activation (**Fig. 9h, 9i, S9e**). It will be important to determine whether this reduction in STING levels in infected primary airway cells is due to direct antagonism by SARS-CoV-2 proteins, such as ORF3a, ORF10 and 3CLpro have been shown to antagonize the cGAS-STING pathway largely through inhibition of downstream signaling ^76–78^. Alternatively, SARS-CoV-2 may preferentially infect of cells that lack STING and hence those cells with lower levels of basal innate immune activation. We note that the effects of STING inhibitors on SARS-CoV-2 replication in in primary cells was relatively modest and that only H-151 reduced innate immune activation whereas SN-011 was ineffective (**Fig. 9h, 9i, S9e**). These discrepancies may in part be due to the inability to fully target STING in the ciliated cells at the air-facing side; STING inhibitors needed to be provided in the liquid phase to maintain ALI integrity with the assumption that they would diffuse to the ciliated cells efficiently and equally. More targeted genetic ablation of STING in primary cells and in vivo models will be needed to address this possible shortcoming.

While our data strongly suggests that cGAS-STING pathway activation underlies the resistance phenotype of multiple ACE2(high) cell models, there were notable differences between these cell lines as well. First, Detroit 562 cells supported only low levels of virus replication upon STING KO which persisted even after ACE2 overexpression, suggesting the presence of other independent blocks to replication. Second, despite successful virus binding/entry (**Fig. 1d, e**) and the notable reduction in numerous IFN pathway genes upon STING and STAT1 KO (**Fig. S3k, 5a**), H596 cells did not support SARS-CoV-2 replication and required ectopic ACE2 expression. Given the importance of TMPRSS2 in defining the cellular tropism of SARS-CoV-2 ^13^, it is important to note that both Detroit 562 and H596 cells expressed comparably higher levels of TMPRSS2 (**Fig. S1a**), suggesting the likelihood of a different post-entry block to replication. Third, pretreatment of SCC25 but not OE21 cells with STING inhibitors was sufficient to enhance SARS-CoV-2 replication, suggesting a higher degree of STING activation in the latter, as also supported by the higher levels of P-IRF3 detected in cell lysates (**Fig. S4b**). Fourth, all cell lines expressed distinct sets of ISGs and inflammatory genes basally, suggesting that the SARS-CoV-2 resistance may be mediated by different effector molecules. Moving forward, understanding the molecular basis of these post-entry blocks through comparative analysis of the cell models presented herein can reveal novel cell intrinsic mechanisms that limit SARS-CoV-2 replication.

IFN signaling leads to the transcriptional upregulation of hundreds of ISGs, many of which are effector molecules that limit virus replication by distinct mechanisms ^79,80^. Several studies have reported on ISG screens and individual antiviral properties of ISGs against SARS-CoV-2, as summarized recently ^2^. Our findings suggest that the ISGs with previously demonstrated antiviral activity against SARS-CoV-2 are unlikely to explain the resistance phenotype of the ACE2(high) cells (**Fig. 4g, 4h**). Whether a dominant ISG or multiple ISGs limit SARS-CoV-2 replication in our cell models remains unknown. Of note, many of the prior ISG screens have been conducted in ACE2 overexpressing cells or cell lines of non-airway origin. In light of our finding that ACE2 overexpression relieves ISG-mediated restriction of SARS-CoV-2 replication (**Fig. 4f, S4f**), we propose that the cell models studied herein provide a unique platform to discover the antiviral activities of ISGs.

SARS-CoV-2 encodes numerous non-structural and accessory proteins that can antagonize IFN-mediated innate immune responses by blocking the production of IFN-β, directly interacting with IFN pathway components and inhibiting IFN signaling, or inhibiting the activities of specific ISGs ^81,82^. Given that ISG expression is limited to uninfected bystander cells in both SCC25 STING KO cells and primary airway cultures, our findings suggest that SARS-CoV-2 replication prevents IFN signaling but not IFN production in infected cells. While these findings are in agreement with a SARS-CoV-2-infected gut organoid system ^83^, others have reported ISG expression in both infected and bystander cells in a similar HBEC-ALI setting ^84^. It remains to be determined whether one or more of the abovementioned SARS-CoV-2 proteins is sufficient to block IFN signaling in the cell models presented here.

Understanding the fundamental biology of how SARS-CoV-2 interacts with the innate immune responses in tractable cell culture-based systems is critical to our understanding of the complex SARS-CoV-2 pathogenesis *in vivo.* Our study highlights that basal activation of the cGAS-STING and IFN pathways can contribute to defining the SARS-CoV-2 cellular tropism at physiologically relevant ACE2 expression levels. While correlative, it is noteworthy that three of the four ACE2(high)/ISG(high) cell lines (SCC25, OE21, Detroit 562) are derived from the upper airways, whereby cGAS-STING mediated restriction may be operative. Our findings also demonstrate the ability of SARS-CoV-2 to efficiently antagonize the IFN pathway, but not RLR-dependent innate immune sensing and IFN production once productive infection is established. Further dissection of these pathways may lead to the identification of actionable host cell targets that are central to the complex SARS-CoV-2 pathogenesis.

## METHODS

Our research complies with all relevant ethical regulations and all relevant protocols have been approved by Washington University School of Medicine Institutional Biological and Chemical Safety Committee. The human airway epithelial cells are isolated from non-living individuals, and therefore research involving the use of these cells is exempt from human subjects regulations as determined by the Washington University Institutional Review Board.

### Cell Culture and Compounds

All cell lines were maintained in a humidified incubator at 37°C with 5% CO_2_ unless otherwise indicated. Cell line identities were validated by short tandem repeat analysis (LabCorp, Genetica Cell Line Testing) and cultures were regularly tested for mycoplasma contamination using the MycoAlert mycoplasma detection kit (Lonza). HEK293T and Calu-3 (ATCC-HTB-55) cells were cultured in Dulbecco’s Modified Eagle’s Medium (DMEM, Sigma), supplemented with 10% FBS (VWR). Vero CCL-81, Vero E6 and Vero E6-TMPRSS2 (kind gift of Whelan lab ^85^) cells were cultured in DMEM, supplemented with 10% FBS and 10 mM HEPES buffer. The A427 lung carcinoma (kind gift from Bernard Weissman Lab (UNC)) and Detroit 562 pharyngeal carcinoma (ATCC-CCL-138) cell lines were cultured in Eagle’s Minimum Essential Medium (EMEM, Corning), supplemented with 10% FBS (Sigma), and 2 mmol/L ʟ-glutamine (Gibco). The SCC25 squamous cell carcinoma (kind gift from the John Hayes Lab (UTHSC)) cell line isolated from tongue was cultured in DMEM:F12 (Corning) supplemented with 10% FBS (Sigma). The H522 non-small cell lung adenocarcinoma (ATCC-CRL-5810), H596 adenosquamous lung carcinoma (ATCC HTB-178), H1299 non-small cell lung carcinoma (ATCC CRL-5803), HCC827 lung adenocarcinoma (ATCC CRL-2868), PC-9 lung adenocarcinoma (Sigma #90071810), KYSE30 oesophageal squamous cell carcinoma (kind gift from the John Hayes Lab), and OE21 oesophageal squamous cell carcinoma (Sigma # 96062201) cell lines were cultured in Roswell Park Memorial Institute (RPMI) 1640 (Corning) with L-glutamine supplemented with 10% FBS (Sigma). THP-1 cells (ATCC#TIB-202) and its derivatives described previously^86^ were grown in RPMI-1640 media supplemented with 10% heat-inactivated fetal bovine serum.

Primary human bronchial epithelial cells (HBECs) grown at air-liquid interface (ALI) were processed as follows. Human airway epithelial cells were isolated from surgical excess of tracheobronchial segments of lungs donated for transplantation as previously described and were exempt from regulation by US Department of Health and Human Services regulation 45 Code of Federal Regulations Part 46 ^87^. Tracheobronchial cells were expanded in culture, seeded on supported membranes (Transwell; Corning, Inc.), and differentiated using ALI conditions as detailed before using 24-well inserts ^88,89^.

The following compounds were reconstituted in solvents following manufacturer’s instructions and stored in aliquots in −80. 2’3’-cGAMP (Invivogen, Cat no: tlrl-nacga23-1), 2’2’-cGAMP (Cayman, Cat no: 22419, H-151 (MedChemExpress, Cat no: HY-112693, Lot no:261833), SN-011 (MedChemExpress, Cat no: HY-145010, Lot no: 149227), VBIT-4, azidothymidine (AZT, NIH AIDS Reagents, Cat no: HRP-3485). Cells were treated as indicated in figure legends and mock-treated cells were included as controls in all experiments.

### Viral Strains

SARS-CoV-2 strain 2019-nCoV/USA-WA1/2020 was obtained from Centers for Disease Control and Prevention (a gift of Natalie Thornburg). The mNeonGreen (mNG) SARS-CoV-2 reporter virus ^90^ was a gift of Pei-Yong Shi lab, and SARS-CoV-2 Lambda, Iota, Mu, Beta, Epsilon, Delta, Kappa, and Omicron variant isolates were obtained from Dr. Michael S. Diamond. SARS-CoV-2, SARS-CoV-2-mNG and variant isolates were propagated in Vero E6-TMPRSS2 at an MOI of 0.01 grown in DMEM (Sigma), supplemented with 2% Fetal Bovine Serum (FBS, VWR) and 10 mM HEPES buffer (Corning). SARS-CoV-2 Omicron variant strain was grown in DMEM supplemented with 1% FBS. After amplification, the viral stocks were titered on Vero-TMPRSS2 cells by plaque assays and sequence confirmed. All experiments involving SARS-CoV-2 were performed in a biosafety level 3 laboratory.

### SARS-CoV-2 infections and focus-forming assay

Cells seeded at 70-80% density were inoculated with virus diluted in cell culture media supplemented with 2% FBS and intermittent rocking for 1 h. Virus inoculum was removed, cells washed with 1x phosphate-buffered saline (PBS) and plated in cell culture media containing 10% FBS. Infections were monitored by focus-forming assay, and RT-qPCR of cell-associated vRNA (N sub-genomic RNA). Briefly, for focus-forming assay, Vero E6-TMPRSS2 cells were inoculated with 10-fold serial dilutions of virus containing supernatants from the panel of airway cell lines, incubated for 1 h at 37°C with intermittent rocking, followed by addition of 2% methylcellulose and 2X MEM containing 4% FBS. 30 hours post-infection cells were fixed by 4% paraformaldehyde (PFA) and stained as previously described ^85^. Briefly, cells were incubated with CR3022 anti-S antibody (1 ug/mL)^91^ overnight followed by anti-human IgG-HRP-conjugated goat secondary antibody (Sigma, 1:500) for 2 h, then foci were visualized with TrueBlue peroxidase substrate (KPL, #5510-0050) and manually counted under the microscope.

Prior to infection, HBECs were washed two times with 1× PBS to remove the mucous layer that otherwise can slow down infection. HBECs were inoculated with SARS-CoV-2-mNG in DMEM-supplemented with 2% FBS for two hours in a humidified incubator at 37°C, after which the initial inoculum was removed, cells washed with 1xPBS and maintained at ALI for the duration of the assays. Subsequently, cells were fixed with 4% paraformaldehyde and subjected to immunofluorescence microscopy as detailed below.

### ACE2 overexpression, lentivirus production and transduction

Cell lines stably expressing human ACE2 were generated by lentiviral transduction as described before^31^. Briefly, lentiviruses were produced in HEK293T cells by polyethyleneimine (Polysciences)-mediated transfection using an pLV-EF1a-IRES-Puro (Addgene Plasmid #85132)-based vector expressing human ACE2, psPAX2 as packaging plasmid (Addgene #12260) and VSV-G expressing envelope plasmid (Addgene #12259). Cell culture supernatants containing lentiviruses were collected and used to transduce SCC25, OE21, Detroit 562, H522 and H596 airway cell lines. Two days post transduction, cells were selected for two weeks in 2 μg/mL puromycin to generate bulk populations expressing ACE2.

### CRISPR knockout cell lines

ACE2 knockout cell lines were generated as previously stated ^31^. In short, lentiviruses were generated in HEK293T cells by transfection of pLentiCRISPRv2 -Puro plasmid containing a single guide RNA (sgRNA) (GTACTGTAGATGGTGCTCAT) (GenScript) targeting ACE2, psPAX2 packaging plasmids (Addgene #12260), and pMD2.G plasmid expressing VSV-G (Addgene #12259) in a ratio of 1:1:0.2 respectively using polyethyleneimine. After transduction of PC-9, KYSE30, HCC827, H1299, SCC25, OE21, A427, Detroit 562, and H596 airway cell lines with the resulting lentiviruses, cells were selected in 2 μg /mL puromycin for two weeks. cGAS-STING pathway (i.e. cGAS, STING, TBK1, and IRF3), STAT1, and TREX1 knockout cell lines were generated similarly by transduction using lentiviral particles generated using pLentiCRISPRv2 -Puro encoding two separate gene-specific sgRNA per target (Table S1), psPAX2, and VSV-G as described above. Following transduction, SCC25, H596, OE21, and Detroit 562 cell lines were selected in 2 μg/mL puromycin for at least two weeks. In all experiments, bulk populations were used for downstream analyses. cGAS and STING KO THP-1 cells were explained before^86^.

### RNA extraction and RT-qPCR

Cell associated RNA was extracted by Trizol following manufacturer’s instructions (Thermo-Fisher Scientific) and was subjected to RT-qPCR analysis for viral (*N* sgRNA) or cellular RNAs. SARS-CoV-2 viral RNA was detected by reverse transcription and amplification using the TaqMan RNA-to-CT 1-Step Kit (Thermo Fisher Scientific), utilizing the primers and probe sequences for viral RNAs are as described before [76]. Briefly, reverse transcription was performed at 48 °C for 15 minutes, followed by a 2-minute incubation at 95 °C. Amplification was carried out over 50 cycles, with each cycle consisting of a 15 second denaturation step at 95 °C, followed by a 1-minute annealing/extension step at 60 °C. The quantification of SARS-CoV-2 N gene RNA copies was determined using a previously published internal RNA standard ^92^. To study the type I/III IFN response at baseline and in response to various stimuli, cellular RNA was reverse transcribed with High-Capacity cDNA Reverse Transcription kit (Thermo Fisher Scientific) followed by RT-qPCR analysis using PowerUp SYBR Green Master Mix (Applied Biosystems), utilizing specific primers that are listed in the Table S2. RNA levels were quantified using the 2 ^-ΔΔCT^ method and normalized relative to *18S rRNA* as the reference target.

### Viral RNA detection by RNAscope

SARS-CoV-2 RNA was detected as described before (41) with minor modifications using the RNAscope Multiplex Fluorescent Reagent Kit v2 (Advanced Cell Diagnostics catalog #323100). One day after attachment on coverslips placed in 24-well plates, cells were infected with SARS-CoV-2 at an MOI of 2 for 1 h, then washed twice with 1x PBS and fresh media was added. Cells were fixed with 4% PFA at 2 hours post infection, dehydrated and stored at −20°C. On the day of detection, cells were rehydrated and incubated in 0.1% Tween in PBS for 10 min and immobilized on slides. Before probing, cells were first treated with hydrogen peroxide (Advanced Cell Diagnostics catalog #322381) and 1:5 diluted protease III (Advanced Cell Diagnostics catalog #322381) each for 10 minutes at room temperature. Immediately after, an anti-sense RNAscope Probe-V-nCoV2019-S (Advanced Cell Diagnostics catalog #848561) targeting SARS-CoV-2 positive strand S RNA was allowed to hybridize at 40°C for 2 hr. All consecutive incubations were performed at 40°C unless otherwise indicated. After probe hybridization, Amplifiers (Amp) were added sequentially as follows; Amp 1 for 30 min, followed by Amp 2 for 30 min, and Amp 3 for 15 min. Samples were then incubated with detection reagents including horseradish peroxidase (HRP)-C1 for 15 min, diluted Tyramide Signal Amplification (TSA) Vivid Fluorophore 520 (1:1500, Advanced Cell Diagnostics catalog #323271) in TSA buffer (Advanced Cell Diagnostics catalog #322809) for 30 min and HRP blocker for 30 min. Nuclei were stained with DAPI at room temperature for 30 sec. Coverslips were mounted on slides using Prolong Gold Antifade. Images were taken using a Zeiss LSM 880 Airyscan confocal microscope equipped with a 63/1.4 oil-immersion objective.

### Immunofluorescence microscopy

SARS-CoV-2 N protein was visualized in infected cells as before ^31^. Briefly, cells grown on coverslips (Neuvitro, #GG-12-1.5-Collagen) were infected with SARS-CoV-2 and fixed with 4% paraformaldehyde. Coverslips were washed with 1X PBS, incubated in 1% bovine serum albumin (BSA) and 10% FBS in PBS containing 0.1% Tween-20 (PBST) at room temperature for 1 h. Samples were then incubated in a primary mouse SARS-CoV-2 nucleocapsid (N) antibody (Sino Biological Inc., Cat # 40588-T62, 1:1000) at 4 °C overnight. After washing with PBST, samples were incubated in a goat anti-mouse secondary antibody conjugated to Alexa Fluor Plus 488 (Invitrogen, Cat# A-11029, 1:1000) at room temperature for 1 h. IFIT3, IFIT2, ISG15 and STING were detected by incubation with a primary rabbit polyclonal IFIT3 antibody (Novus Biologicals NBP2-32500, 1:500), rabbit polyclonal IFIT2 antibody (Novus Biologicals NBP2-15180SS, 1:500), rabbit polyclonal ISG15 antibody (Proteintech 15981-1-AP, 1:250) and rabbit polyclonal STING antibody (Proteintech 19851-1-AP, 1:200) respectively as described above followed by incubation with a goat anti-rabbit fluorescent secondary antibody (Invitrogen Goat anti-Rabbit Alexa Fluor Plus 568™, Cat# A-11036, 1:1000 or Invitrogen Goat anti-Rabbit Alexa Fluor™ Plus 647, Catalog # A32733TR, 1:1000). Nuclei were stained with DAPI diluted in PBS at room temperature for 5 min. Finally, coverslips were washed in PBST followed by PBS and then mounted on slides using Prolong Gold Antifade.

For confocal microscopy, samples were analyzed with a Zeiss LSM880 laser scanning confocal microscope (Carl Zeiss Inc. Thornwood, NY). The system is equipped with 405nm diode, 488nm Argon, 543nm HeNe, and 633nm HeNe lasers. Plan-Apochromat 20X numerical aperture (NA) 0.8), Plan-Apochromat 40X (NA 1.4) DIC oil, and Plan-Apochromat 63X (NA 1.4) DIC oil objectives were used. ZEN black software (version 2.1 SP3) was used for multichannel image acquisition and for obtaining Z stacks with optimal interval and section thickness. The image analysis software Volocity with Visualization module (version 6.3) (PerkinElmer, Waltham, MA) was used for 3-dimensional rendering of Z slices acquired through the depth of the cell monolayers. Colocalization analysis of Z stacks was performed using the Colocalization module of Volocity. For epifluorescence microscopy, images were taken using Leica using either the ×4 or ×20 objective.

### Immunoblotting

Human cell lines were grown to 70% confluence and lysed in 1X RIPA buffer (50mM Tris-HCl pH 7.4, 150mM NaCl, 1mM EDTA, 0.1% SDS, 1% NP-40, 0.25% sodium deoxycholate) supplemented with EDTA-free protease inhibitor cocktail (Roche). Proteins were separated by SDS-PAGE on 4-12% Bis-Tris gels, transferred to a nitrocellulose membranes, blocked in Intercept (PBS) blocking buffer (LI-COR) or 5% milk, and incubated with primary antibodies overnight at 4°C. Washed membranes were incubated for 45 min to 1 hour at room temperature in secondary antibody solution (LI-COR IRDye 680, 800; 1:10,000 in 5% milk or Intercept (PBS) blocking buffer), imaged on an Odyssey® CLx scanner, and analyzed using the Image Studio Software. Antibodies were used at the following dilutions: ACE2 (R&D Systems #AF933, 1:200), β-actin (Sigma #A5316, 1:5000 or Santa Cruz Biotech #sc8432, 1:500), Vinculin (Santa Cruz #sc-73614, 1:2000), pSTAT1-Y701 (Cell Signaling #9167, 1:1000), pSTAT1-S727 (Cell Signaling #8826, 1:1000), STAT1 (Cell Signaling #14994, 1:1000), MX1 (Cell Signaling #37849, 1:1000), IFIT1 (Cell Signaling #14769, 1:1000), STING (Cell Signaling #13647S, 1:1000), cGAS (Cell Signaling #15102S, 1:1000), TBK1 (Cell Signaling #38066S, 1:1000), IRF3 (Cell Signaling #4302S, 1:1000), SARS-CoV-2 Nucleocapsid (Sino Biological # 40588-T62, 1;1000 or # 40143-MM05, 1:1000), P-STING S366 (Cell Signaling #D7C3S, 1:1000), P-IRF3 S396 (Cell Signaling #4D4G, 1:1000) and TREX1 (Cell Signaling #15107S, 1:1000) antibodies.

### Flow cytometry

SARS-CoV-2 mNG infected cells were detached with trypsin and fixed with 4% paraformaldehyde at room temperature for 20 min. After fixation, cells were collected by centrifugation, resuspended in 1X PBS and subjected to flow cytometry directly or processed for staining. For staining, cells were permabilized in 1X PBS supplemented with 0.5% Tween-20 (PBST) for 10 min, washed once with 1X PBS and blocked by incubation in 1X PBST supplemented with 1% bovine serum albumin and 10% FBS for 1h at room tempreature. Samples were then incubated in a primary rabbit polyclonal antibody against IFIT3 (Novus Biologicals, NBP2-32500, 1:500) at 4 ° C overnight followed by staining with a goat anti-rabbit Alexa Fluor Plus 647 (Life Technologies, A32733TR, 1:1000) for 1 h at room temperature. Cells were washed in 1X PBST in between antibody incubations and resuspended in 1X PBS. Flow cytometry was performed using BD LSR Fortessa flow cytometer and analyzed by FlowJo software.

### siRNA transfections

siRNAs (Table S3) were reverse-transfected as described before ^31^. In brief, 5 pmoles of siRNAs for each target was complexed with 1.5 µL of Lipofectamine RNAiMAX transfection reagent in 50 μL of Opti-MEM following manufacturer’s recommendations and added in 24-well cell culture dishes. SCC25 or OE21 cells were then seeded at 1×10^5^ cells/well. At 2-days post-transfection cells were infected with SARS-CoV-2 at an MOI of 2 i.u./cell, and RNA extracted 3 days later by Trizol and processed for RT-qPCR analysis as above. Knockdown efficiency was assessed by RT-qPCR or immunoblotting.

### Analyses of IFN activity in culture supernatants and secondary messengers in the cytosol

To determine whether SCC25, H596, OE21 and Detroit 562 cells actively release type I/III IFNs in cell culture supernatants compared to other airway-derived cell lines, 5×10^5^ cells were plated in 6-well dishes. At 24h post-plating, culture supernatants were collected, centrifuged at 500x*g* to remove cell debris and 500 μl of the resultant supernatant was inoculated on 2×10^5^ THP-1 cells (or its derivatives thereof, plated in 500 μl RPMI) in 24-well plates. Total RNA was isolated at 24h post-inoculation by Trizol and subjected to qRT-PCR as detailed above. In a parallel set of experiments, 5×10^5^ cells were plated in 6-well dishes one day prior to co-culturing. On the day of co-culturing, spent cell culture medium was removed, cells washed with PBS to remove cell debris and unattached cells and 1×10^6^ THP-1 cells added on top. 24h post co-culturing, THP-1 cells that remain in suspension were gently removed and subjected to RNA extraction and qRT-PCR. Attachment properties of airway-derived cells were visually assessed by microscopy and we found no major impact in cell morphology or viability following co-culturing.

To assess the presence of cGAS-derived secondary messengers in airway-derived cells, THP-1 and THP-1 cGAS KO and THP-1 STING KO cells were inoculated with post-nuclear supernatants isolated as follows. In brief, airway-derived cells grown at confluency in 10-cm culture dishes were trypsinized, collected in culture media, washed with 1xPBS and resuspended in 1 ml of hypotonic buffer (10 mM Tris-Cl pH 8.0, 10 mM KCl, 1 mM EDTA supplemented with complete protease inhibitors (Roche)). Following 10 min incubation on ice, cells were dounce-homogenized for 40 strokes and lysates were centrifuged at 1000 x *g* at 4°C for 5 min. 150 μl of resultant PNS was added on 2×10^5^ THP-1 cells in 24-well plates (alongside with controls including 2’3’-cGAMP and 2’2’-cGAMP) and IFN-β expression analyzed by qRT-PCR following RNA extraction by Trizol as detailed above.

Cell culture supernatants were also subjected to an IFN-β ELISA assay (InvivoGen, #luex-hinfbv2) or IFN-γ ELISA assay (DY1598B-05 from R&D systems) following the manufacturer’s instructions. The concentration of IFNs in cell culture supernatants was derived based on a standard curve obtained from serial dilutions of recombinant IFNs provided by the ELISA assay.

### Analysis of cytosolic mtDNA

Presence of cytosolic mtDNA in the cell line panel was analyzed as described before^93^. In brief, cells grown to near confluency in 10-cm dishes were collected by trypsin, washed with 1xPBS and sequentially lysed in 500 μl of buffers containing digitonin, NP-40 and SDS that allow permeabilization of the plasma membrane, cellular membranes including mitochondrial and nuclear membrane, respectively. 50 μl of the extract was proteinase K treated and DNA extracted by phenol:chloroform:isoamyl alcohol extraction. Resultant DNA was analyzed by qPCR using KCNJ as the nuclear marker and MT-ND1 and MT-D-loop as mitochondrial markers. Abundance of mtDNA in the cytosol was normalized relative to its total abundance in each cell type as described in detail before^93^.

### Gene expression analysis in infected vs. bystander SCC25 cells

1×10^7^ SCC25-STING KO cells were infected with SARS-CoV-2 mNG at MOI: 1 i.u./cell for 72 hrs. Cells were subsequently fixed with 4% paraformaldehyde and single-cell sorted on a BD FACS Aria III platform. Infected (mNG+) and bystander (mNG-) cells were lysed in 1X Proteinase K buffer (100 mM Tris-pH7.5, 50 mM NaCl, 10 mM EDTA, 1% SDS) supplemented with 20 μg of Proteinase K by incubation at 55 °C for 1h by constant agitation on a thermal mixer. RNA was extracted by phenol:chloroform:isoamyl alcohol and pelleted by ammonium acetate and ethanol. Pelleted RNA was subjected to qRT-PCR as above.

### RNA-seq and single-cell RNA-seq data analysis

For heatmap analysis of the interferon response pathway in our cell line cohort, RNA-seq TPM data for the Cancer Cell Line Encyclopedia (CCLE) was downloaded from the DepMap portal (DepMap Public 18Q3). Gene sets for the interferon response pathway were extracted from MsigDB’s lists of HALLMARK_INTERFERON_GAMMA_RESPONSE and HALLMARK_INTERFERON_ALPHA_RESPONSE genes. Hierarchical clustering and figure generation was performed with the ComplexHeatmap R package. For single-cell expression levels in healthy human airways, normalized counts and cell type metadata were downloaded from a publicly available single-cell study ^94^.

For RNA-seq analysis of SCC25, H596, OE21, Detroit 562 cell lines and their STING CRISPR derivatives, total RNA was harvested from cells grown on 6-well plates by Trizol (Life Technologies) following manufacturer’s instructions. 4 μg of total RNA (from n=4 replicates) was subjected to Illumina stranded mRNA library generation kit and sequenced on an Illumina Nextseq 500 platform at 1×75. Resultant RNA-seq reads were aligned to human genome (hg19) using the STAR aligner (add ref) with a mismatch rate threshold of 4% (--outFilterMismatchNoverLmax 0.04). “featureCounts”, a read summarization program, was then used to annotate the mapped reads. RNA-seq counts generated by featureCounts were processed using the edgeR package. The data was filtered with “filterByExpr” with default settings, normalized, and transformed to counts per million (CPM). Genes with CPM values greater than 1 were kept, and their CPM values were log10 transformed. These log10-transformed CPM values were then used to perform PCA using “prcomp” function in R. RNA-seq data on WT and STING knockout airway cells have been deposited to the GEO Server under accession number GSE271679.

The “exactTest” function from edgeR was used to test for genewise differential expression between WT and KO samples of each cell line using log2 CPM values, yielding logFC, P-value and FDR. For the full heatmap presented in Fig. S5, both upregulated and downregulated genes (|logFC| > 1) with FDR < 0.05 were considered. Z-scores of log2-transformed CPM values for all selected genes were calculated and used for consensus clustering with the “ConsensusClusterPlus” package. The “ComplexHeatmap” package was then used to generate the heatmap.

For the heatmap of selected genes in Fig. 5, only those from the human gene sets

“HALLMARK_INFLAMMATORY_RESPONSE,”

“HALLMARK_INTERFERON_ALPHA_RESPONSE,” and

“HALLMARK_INTERFERON_GAMMA_RESPONSE” retrieved from Gene Set Enrichment Analysis (GSEA) were included. Differential expression results were filtered to keep only downregulated genes (logFC < −1) and those with FDR < 0.05.

#### Enrichment Heatmap for gene sets

Overrepresentation analysis was performed with the enricher function from the “ClusterProfiler” package, using CPM values obtained from previous steps. Gene sets that were retrieved from Molecular Signatures Database (MSigDB) of GSEA were selected manually and plotted using the “ComplexHeatmap” package. For enricher function, a cutoff of q < 0.01 was used to define overrepresented gene sets, the minimum number of genes in the set was 10 and the p-value adjustments for multiple comparisons were set to be “fdr”. The −log10-transformed q-values were then calculated and used for plotting the enrichment heatmap.

#### Variant Calling

VCF files were generated using BCFtools with mapped reads. SNPs (single nucleotide polymorphisms) from the VCF files were filtered using the “GATK” package based on the following criteria: QUAL ≥ 30.0, DP ≥ 10.0, QD ≥ 2.0, and SOR ≤ 4.0. Reads that failed to meet these criteria were filtered out. SnpEff was then used to annotate the remaining SNPs.

### Statistical Analysis

Generally, experiments were repeated with at least two biological replicates, represented by n. Each plot includes points for individual biological replicates and mean ± SEM error bars unless otherwise specified. GraphPad Prism 9 software was used for statistical analysis. Statistical parameters and details for each experiment are reported in respective figure legends.

## DATA AVAILABILITY STATEMENT

RNA-seq data on WT and STING knockout airway cells have been deposited to the Gene Expression Omnibus Database under accession number GSE271679 and are available at the following URL: https://www.ncbi.nlm.nih.gov/geo/query/acc.cgi?acc=GSE271679. Source data are provided with this paper.

## CODE AVAILABILITY STATEMENT

Processing of the sequencing data are done by open access software and thoroughly explained in the Methods section. Any additional information required to reanalyze the data reported in this paper is available from the lead contact upon request.

## ACKNOWLEDGEMENTS

This work was supported in part by a V Foundation grant (T2014-009) to M.B. Major, a T32 training grant (T32CA009547-34) to K.M.L., a F31 grant to J.E.E. (AI167695). We thank the Miner lab for STING CRISPR plasmid and the Molecular Microbiology Imaging Facility (Wandy L. Beatty) for help with confocal microscopy and data analysis. We thank members of the Lopez, Lenschow, Diamond, Orvedahl and Boon labs for reagents and critical input. We opted to cite recent review articles where appropriate and apologize to colleagues whose work we were unable to cite due to citation limits.

## AUTHOR CONTRIBUTIONS

M. P-C., K.M.L., M.B.M., D.G. and S.B.K. conceptualized the study; M. P-C., J.E.E., K.M.L., S.D., M.B.M., D.G., S.L.B. and S.B.K. designed the methodology. M. P-C., J.E.E., M.X., K.M.L., J.P., J.X., Z.Z.., S.M., Q.W., D.Q.L., S.D., G.H. and S.B.K performed the experiments. Q.Z., R.J. and D.G. performed statistical and bioinformatics analysis. M. P-C., M.B.M., D.G. and S.B.K. wrote the manuscript with input from all the authors.

