## Supplemental Files for "A basally active cGAS-STING pathway limits SARS-CoV-2 replication in a subset of ACE2 positive airway cell models"

**Running title: Basally active cGAS-STING limits SARS-CoV-2 infection**

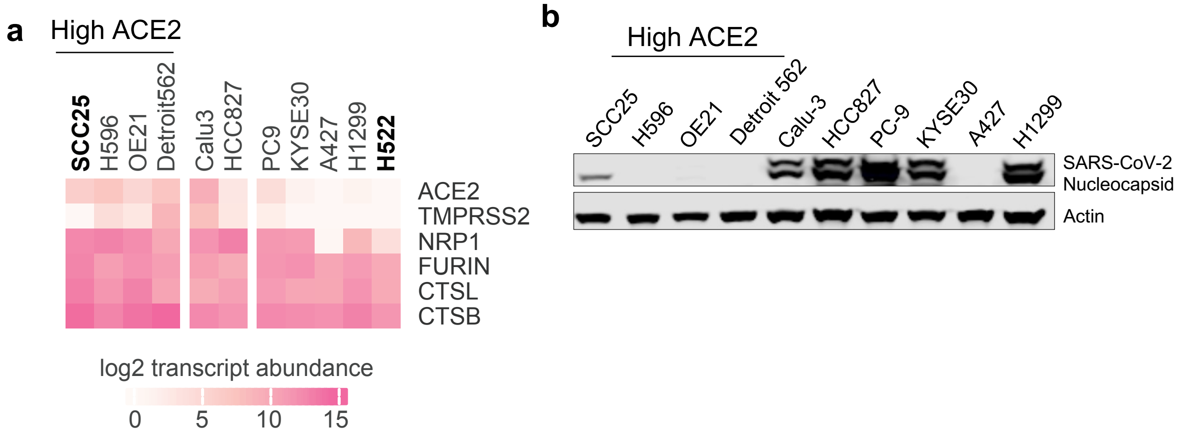


**Figure S1: Analysis of SARS-CoV-2 replication in a panel of airway-derived cell lines that endogenously express ACE2.**

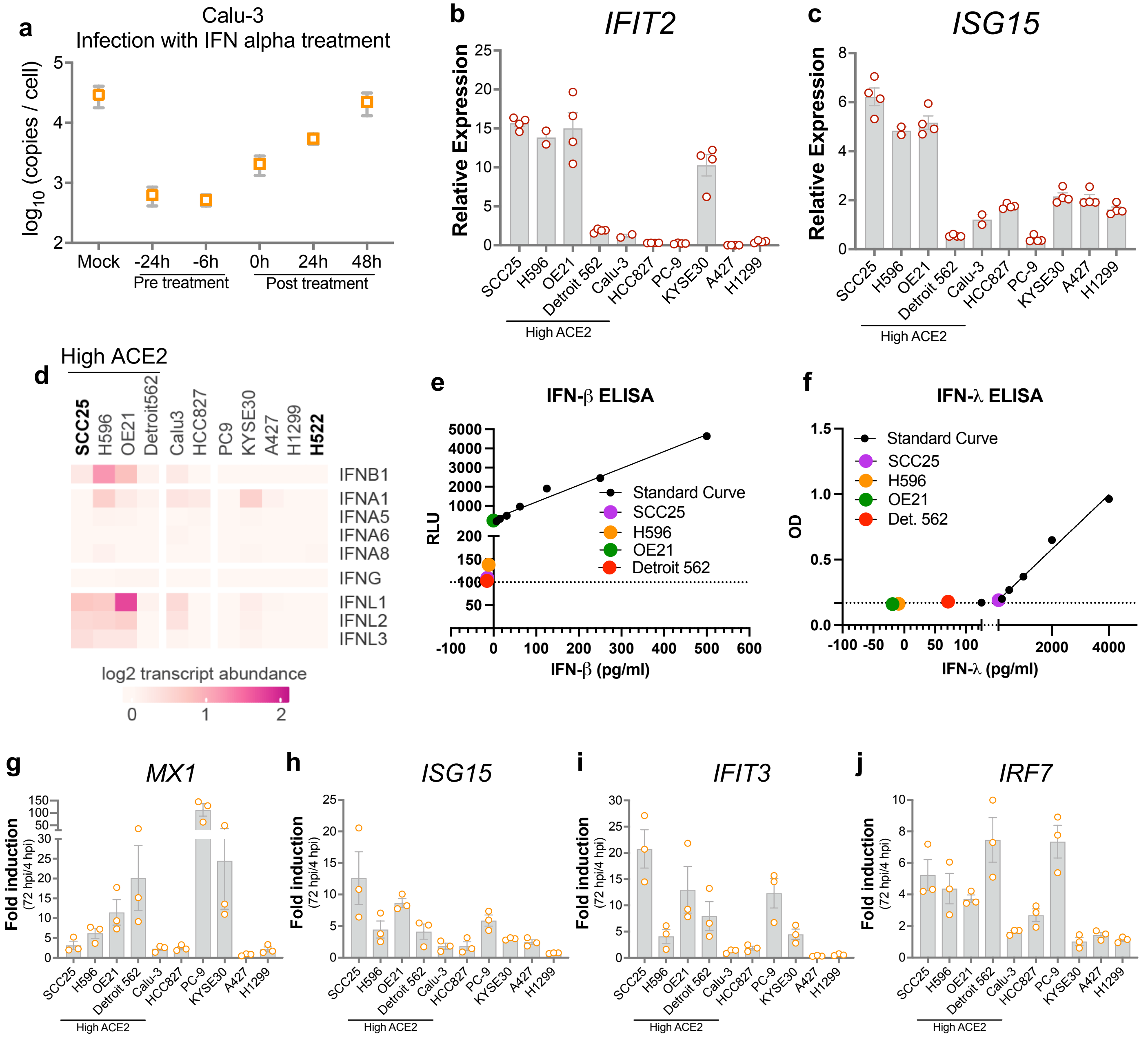


**Figure S2: Expression levels of IFN pathway genes in uninfected and infected cell lines.**

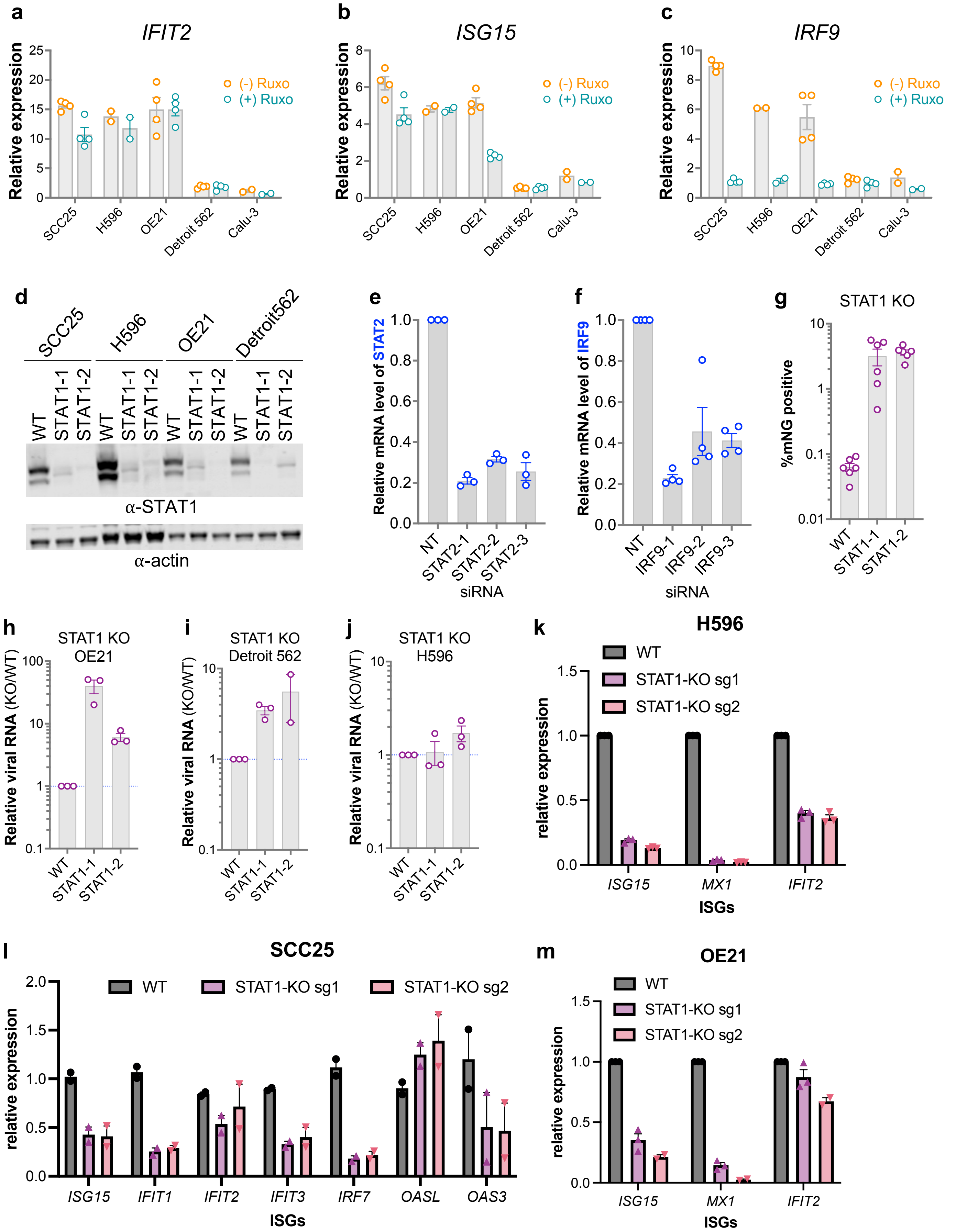


**Figure S3. Basally active type I IFN pathway limits SARS-CoV-2 replication in a subset of ACE2(high) cells.**

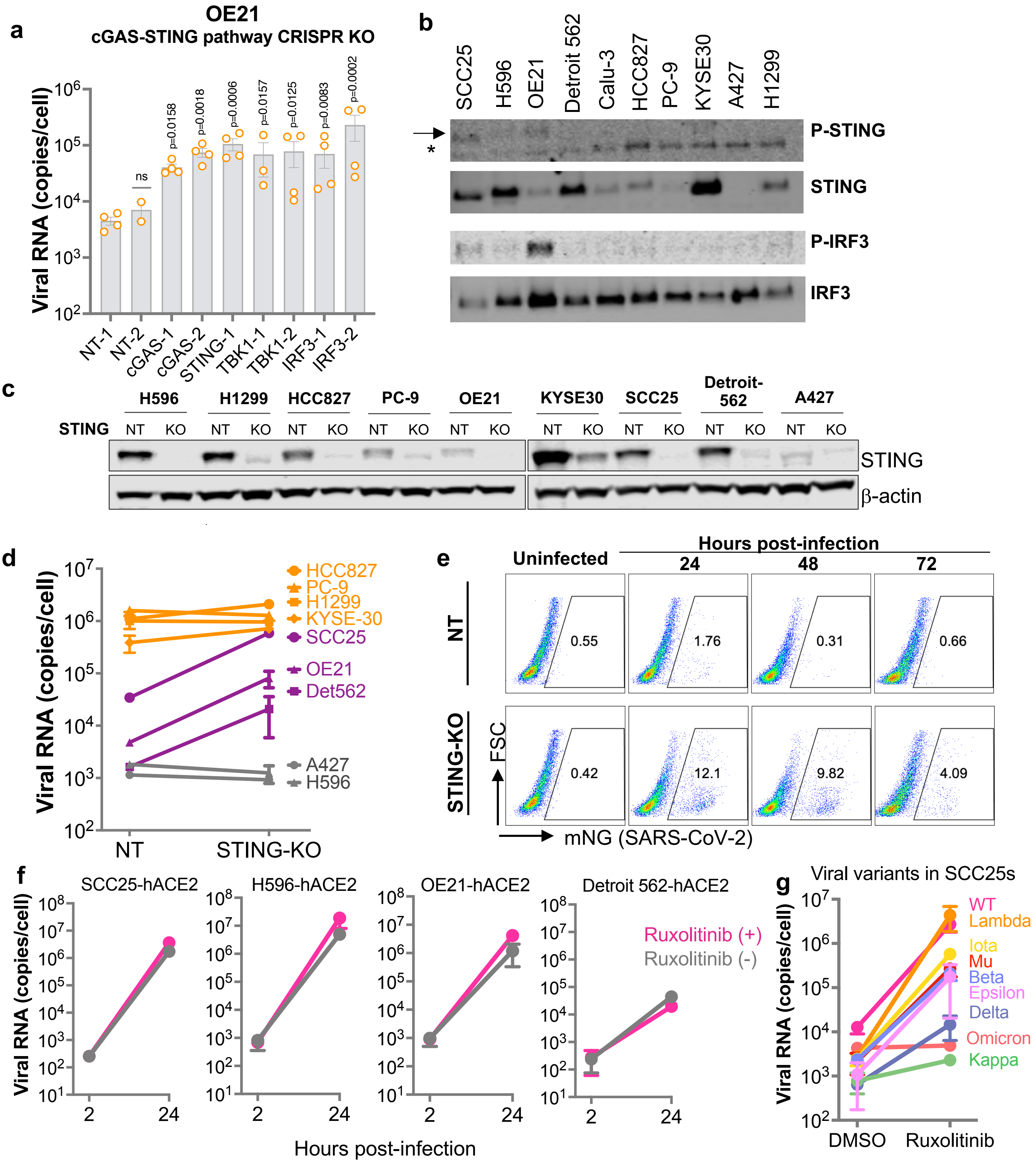


**Figure S4. A constitutively active cGAS-STING pathway limits SARS-CoV-2 replication in a subset of ACE2(high) airways cell lines.**

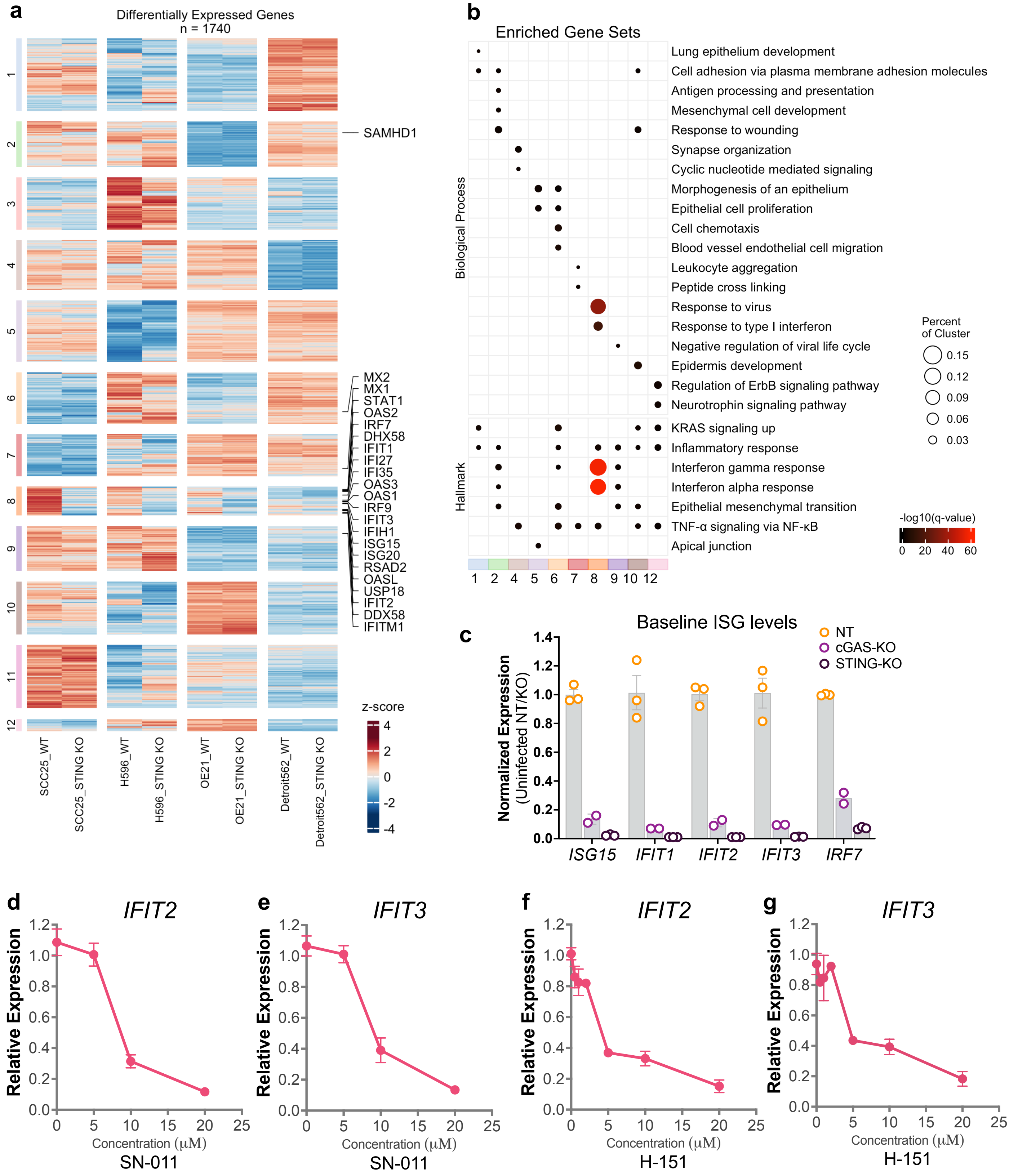


**Figure S5. Genetic ablation and chemical inhibition of STING reduces basal ISG levels and enhances SARS-CoV-2 replication in SCC25 cells.**

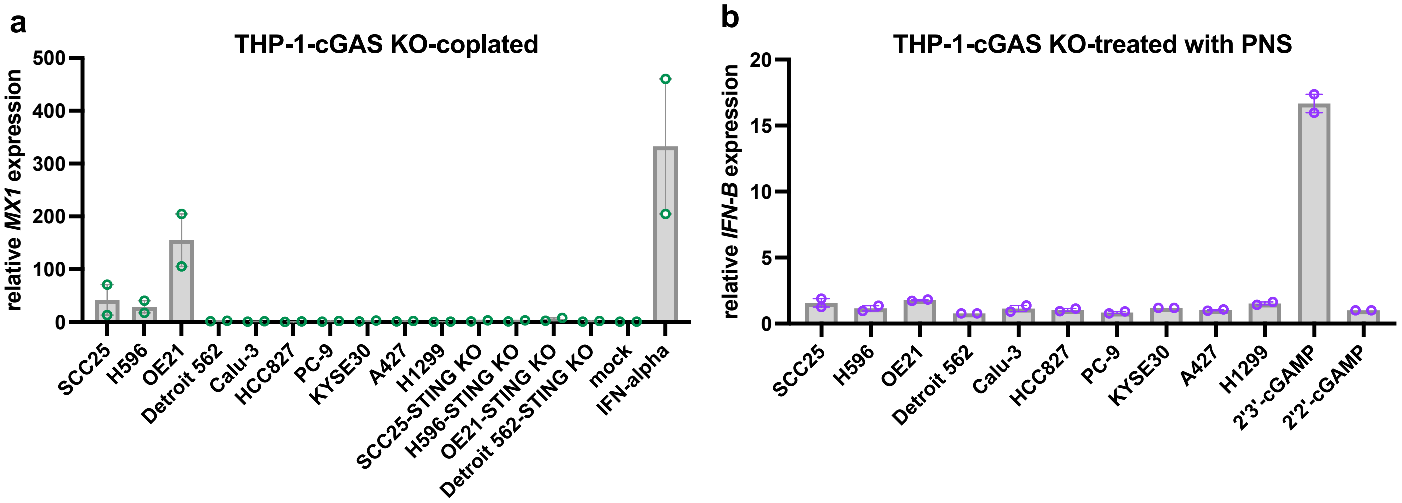


**Figure S6. cGAS-dependence of innate immune activation in target cells co-cultured with and exposed to lysates of ISG(high) cells.**

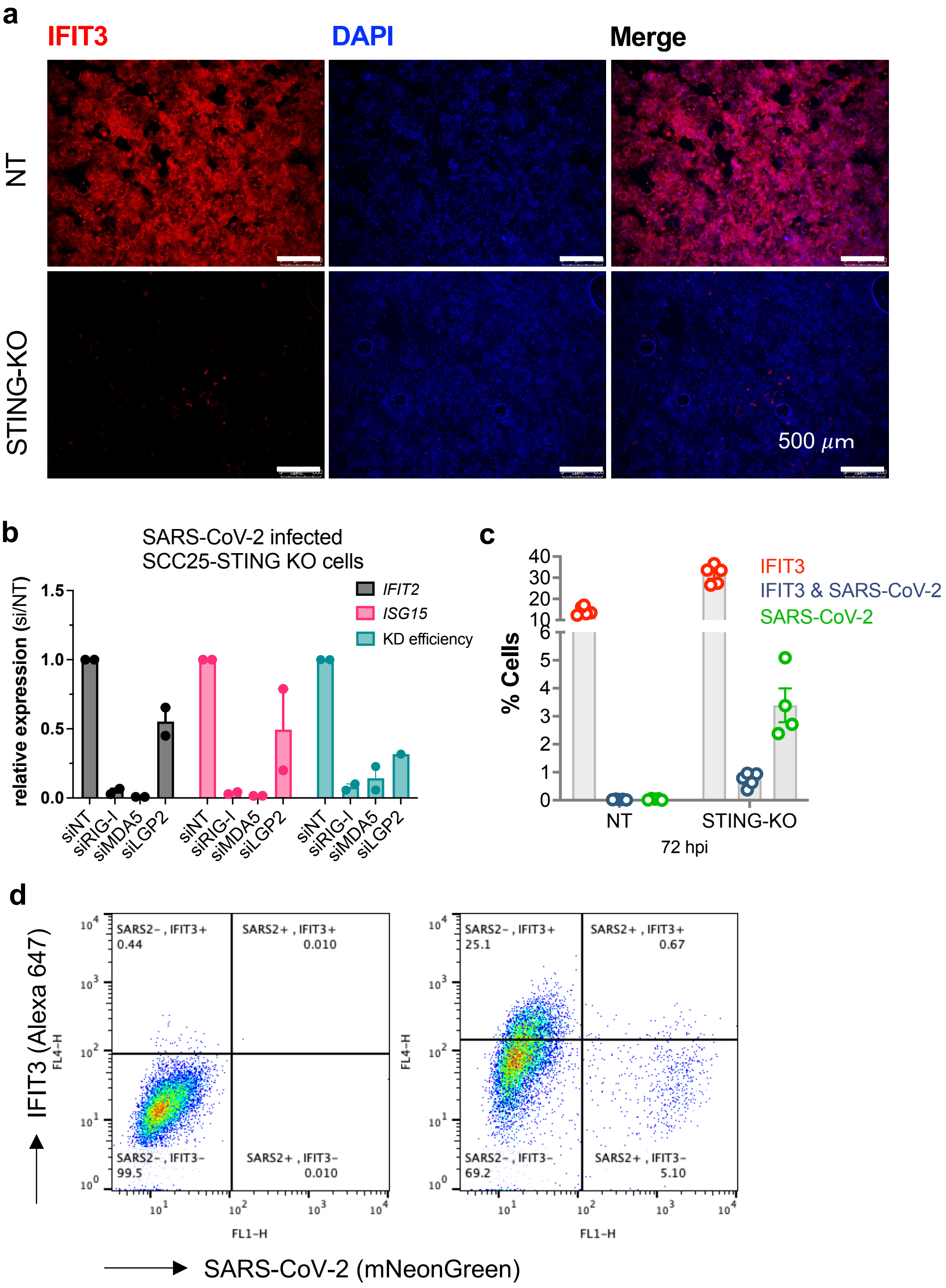


**Figure S7. Impact of an active cGAS - STING pathway-mediated vs. SARS-CoV-2-induced type-I IFN activation on SARS-CoV-2 replication.**

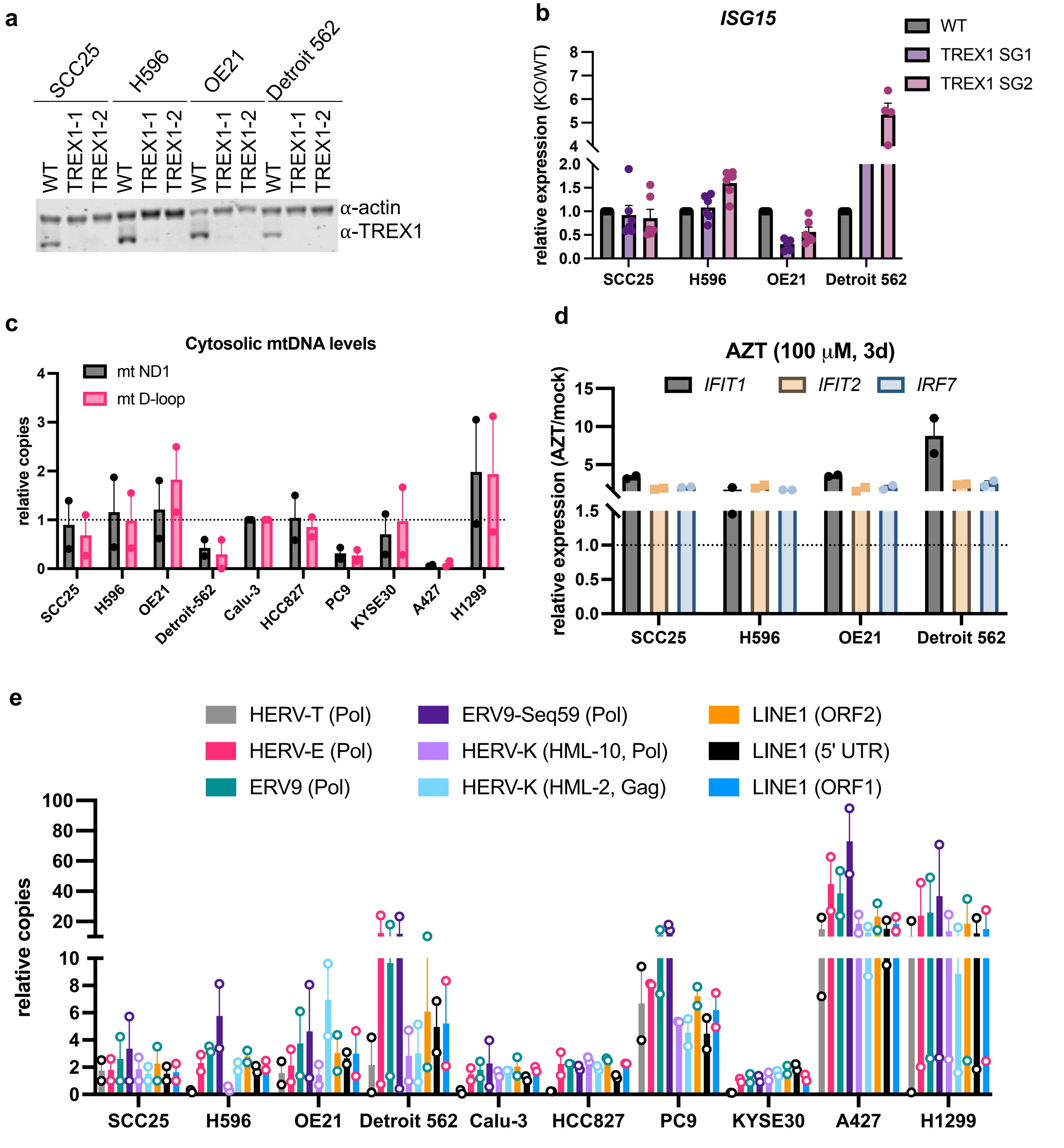


**Figure S8. Analysis of mechanisms that underlie tonic cGAS-STING activation in the ISG(high) cell lines.**

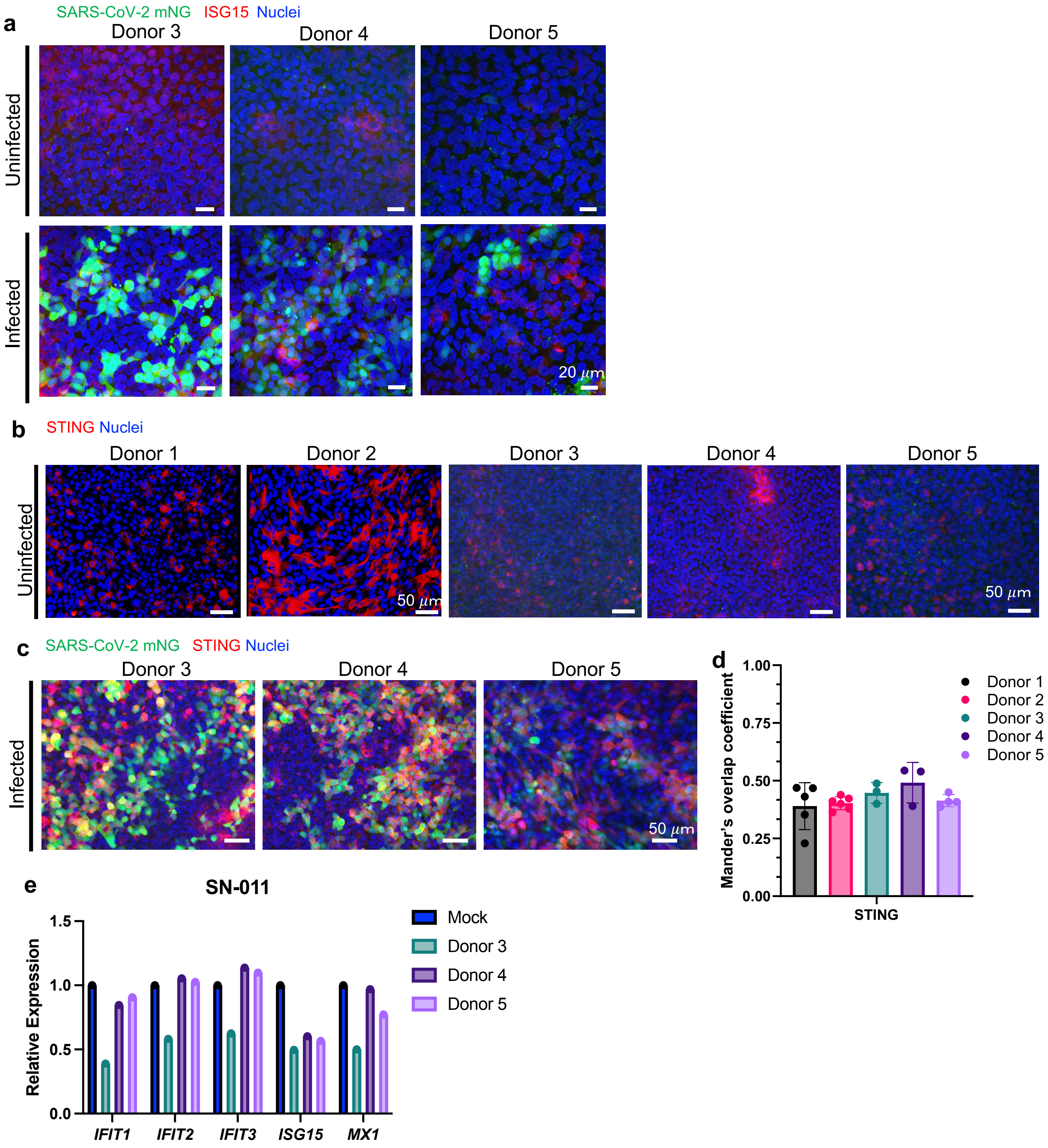


**Figure S9. Analysis of ISG and STING expression in SARS-CoV-2-infected primary human airway epithelial cells (hTECs) grown at ALI.**

### SUPPLEMENTARY FIGURE LEGENDS

**Figure S1: Analysis of SARS-CoV-2 replication in a panel of airway-derived cell lines that endogenously express ACE2. (a)** RNA expression of SARS-CoV-2 viral entry factors downloaded from the Cancer Cell Line Encyclopedia (CCLE). The heatmap displays $log_2$(TPM) values. **(b)** Airway-derived cell lines were infected at a MOI: 2 i.u./cell and cell lysates were analyzed by immunoblotting using anti-SARS-CoV-2 N and anti-actin antibodies at 3 dpi. Immunoblots are representative of two independent replicates.

**Figure S2: Expression levels of IFN pathway genes in uninfected and infected cell lines. (a)** Calu-3 cells were pretreated for 6 or 24 hours or treated at the indicated time points post-infection with 1000 U/mL IFN-$\alpha$ and infected with SARS-CoV-2 at MOI: 2 i.u./cell. RT-qPCR analysis of cell-associated SARS-CoV-2 N sgRNA (copies/cell) at 72 hpi is shown. Data show the average of 4 biological replicates, errors bars denote the SD. **(b, c)** RT-qPCR analysis of representative ISGs including *IFIT2 and ISG15* in uninfected cells. Expression levels are normalized relative to Calu-3 cells (set to 1). Data show the mean from n=2-4 independent biological replicates, error bars denote SEM. **(d)** RNA expression of various IFNs and subtypes was downloaded from the Cancer Cell Line Encyclopedia (CCLE). The heatmap displays $log_2$(TPM) values. (**e, f**) Cell culture supernatants from indicated cell lines were assayed for IFN-$\beta$ (e) or IFN-$\lambda$ (f) using ELISA as detailed in Methods. **(g-i)** The indicated airway-derived cell lines were infected with SARS-CoV-2 at a MOI: 2 i.u./cell and expression of ISGs including *MX1, ISG15, IFIT3 and IRF7* was analyzed by RT-qPCR at 4 and 72 hpi. Y-axes indicate ISG fold induction at 72 hpi relative to 4hpi. Data show the mean from n=3 independent replicates, errors bars show the SEM.

**Figure S3. Basally active type I IFN pathway limits SARS-CoV-2 replication in a subset of ACE2(high) cells. (a-c)** SCC25, H596, OE21, Detroit 562 and Calu-3 cells were pretreated with

1 μM ruxolitinib for 24 hours. *IFIT2* (**a**), *ISG15* (**b**) and *IRF9* (**c**) expression was analyzed by RT-qPCR and expression levels are shown relative to untreated Calu-3 cells (set to 1). Data show the mean from n=2-4 independent biological replicates, error bars show SEM. **(d)** Polyclonal populations of SCC25, H596, OE21 and Detroit 562 cells knocked out (KO) for or STAT1 using two independent sgRNAs were analyzed alongside with parental wild-type (WT) controls for STAT1 and actin by immunoblotting (representative of n=2). **(e-f)** Expression levels of *STAT2* (**e**) and *IRF9* (**f**) were assessed by RT-qPCR in SCC25 cells that were transfected with 3 separate siRNAs targeting STAT2 and IRF9 respectively or a non-targeting control (NT) and infected with SARS-CoV-2 as in Fig. 3. Data display the mean of 3-4 biological replicates, error bars show SEM. (**g**) SCC25 STAT1 KO cells as well as parental WT controls were infected with SARS-CoV-2 mNG at MOI: 2 i.u./cell and percentage of infected cells (mNG positive) was enumerated by flow cytometry at 72 hpi and represented as percentage of total population. Data display the mean of 6 replicates, error bars show SEM. (**h-j**) Analysis of cell-associated SARS-CoV-2 N RNA at 72 hpi (MOI: 2 i.u./cell) in WT and STAT1 KO OE21 (**h**), Detroit 562 (**i**) and H596 (**j**) cells. Data show the mean from 2-3 independent biological replicates, error bars show the SEM. (**k-m**) RT-qPCR analysis of the indicated ISGs in uninfected WT and STAT1 KO H596 (**k**), SCC25 (**l**) and OE21 (**m**) cells. Data display the mean of 3 independent biological replicates, error bars show the SEM.

**Figure S4. A constitutively active cGAS-STING pathway limits SARS-CoV-2 replication in a subset of ACE2(high) airways cell lines. (a)** Polyclonal populations of OE21 cells knocked out (KO) for cGAS, STING, TBK1 and IRF3 using two separate sgRNAs and the non-targeting (NT) controls were infected with WT SARS-CoV-2 at MOI: 2 i.u./cell. RT-qPCR analysis of cell-associated viral N RNA shows copies per cell from 4 independent experiments where error bars show SEM. Indicated p values are derived from one-way ANOVA and the Dunnett correction for multiple comparisons. **(b)** Immunoblots showing total and phosphorylated STING and IRF-3 (P-STING, P-IRF3) in cell lysates. Blots are representative of two replicates. (*) indicates a nonspecific band. **(c)** STING immunoblot of the panel of airway-derived STING KO or NT control cell lines. β-actin serves as the loading control. Immunoblots are representative of two independent replicates. **(d)** Panel of airway-derived STING KO cell lines or the NT controls were infected with SARS-CoV-2 at a MOI: 2 i.u./cell. RT-qPCR analysis of cell-associated viral RNA (copies/cell) at 72 hpi in NT vs. STING KO cells is displayed. Data show the average of 2 biological replicates, where errors bars indicate the SEM. (**e**) Bulk populations of SCC25 STING KO and NT control cells were infected with SARS-CoV-2-mNG at MOI:2 i.u./cell. Percentage of NG-positive cells was determined by flow cytometry at 24, 48 and 72 hpi. Plots are representative of n=2 independent replicates. (**f**) Airway-derived cell lines ectopically expressing ACE2 were pretreated with 1 μM ruxolitinib or mock-treated for 24 hours and infected with SARS-CoV-2 at a MOI: 0.1 i.u./cell in the absence and presence of 1 μM ruxolitinib. RT-qPCR for cell-associated SARS-CoV-2 RNA at 2 and 24 hpi is shown from n=2 replicates. Data show the mean, error bars denote SEM. (**g**) SCC25 cells were pretreated with 1 μM ruxolitinib or DMSO for 24 hours and infected with SARS-CoV-2 Variants of Concern (VOC) at a MOI: 2 i.u./cell in the presence and absence of ruxolitinib. RT-qPCR analysis shows cell-associated viral RNA levels at 72 hpi. Data show the mean from n=2 independent experiments, error bars show SEM.

**Figure S5. Genetic ablation and chemical inhibition of STING reduces basal ISG levels and enhances SARS-CoV-2 replication. (a)** WT and STING KO SCC25, H596, OE21 and Detroit 562 cells were subjected to RNA-seq analysis. Heatmap shows differentially expressed genes (absolute $\log_2$ fold change of >2 and an adjusted *q* value of <0.05 from n=4 independent replicates for each cell type). (**b**) Hypergeometric enrichment analysis from Hallmark and Gene Ontology databases for each individual cluster in A. Color represents significance (*q* value); size indicates the percentage of the cluster represented in the pathway. (**c**) Expression levels of ISGs *ISG15, IFIT1, IFIT2, IFIT3 and IRF7* were assessed by RT-qPCR in uninfected polyclonal populations of SCC25 cells knocked out (KO) for cGAS and STING or transduced with a non-targeting (NT) control. Expression levels were normalized relative to NT control (set to 1). Data show the mean from n=2-3 independent biological replicates, error bars show SEM. (**d-g**) RT-qPCR analysis of *IFIT2* and *IFIT3* in uninfected SCC25 cells treated with increasing concentrations of SN-011 (**d, e**) and H-151 (**f, g**) for 48 h.

**Figure S6. cGAS-dependence of innate immune activation in target cells co-cultured with and exposed to lysates of ISG(high) cells. (a)** *MX1* expression was analyzed by RT-qPCR in THP-1 cGAS KO cells co-cultured with the indicated cell line panel as explained in Methods. Cells treated with 1000 u/ml IFN-$\alpha$ served as the positive control. Data show the mean induction relative to cells without co-culturing (mock) from two independent biological replicates, error bars show the SEM. (**b**) THP-1 cGAS KO cells were treated with post-nuclear supernatants (PNS) from the indicated cells, 10 $\mu$g/ml 2'3'-cGAMP or 10 $\mu$g/ml 2'2'-cGAMP for 24 h, as explained in Methods. IFN-$\beta$ expression was analyzed by RT-qPCR and normalized relative to 2'2'-cGAMP-treated samples. Data show the mean from n=2 independent experiments, error bars show the SEM.

**Figure S7. Impact of an active cGAS - STING pathway-mediated vs. SARS-CoV-2-induced type-I IFN activation on SARS-CoV-2 replication.** (**a**) Uninfected polyclonal populations of STING KO or NT SCC25 cells were subjected to immunostaining for IFIT3. IFIT3 label (in red) and nuclei (DAPI, in blue) observed under 4X objective by epifluorescence microscopy. Scale bars show 500 $\mu$M, (representative of n=2). (**b**) SCC25 STING KO cells were transfected with siRNAs targeting RIG-I, MDA-5 and LGP-2 or a non-targeting (NT) control as explained in Methods. Two days post-transfection, cells were infected with WT SARS-CoV-2 at MOI: 2 i.u./cell for 72 h. *IFIT2* and *ISG15* expression as well as knockdown efficiency was analyzed by RT-qPCR and normalized relative to NT samples (set to 1). Data show the mean from two independent replicates, error bars show the SEM. **(c)** Bulk populations of SCC25 STING KO cells were infected with SARS-CoV-2 at a MOI:2 i.u./cell. Percentage of SARS-CoV-2 nucleocapsid (NP)-positive cells (green) and IFIT3 (red) was determined by flow cytometry at 72 hpi. Data show the mean of 4 biological replicates, error bars denote the SEM. (**d**) Exemplary FACS plots of data shown in Panel (c).

**Figure S8. Analysis of mechanisms that underlie tonic cGAS-STING activation in the ISG(high) cell lines. (a, b)** Polyclonal populations of SCC25, H596, OE21 and Detroit 562 cells knocked out (KO) for TREX1 using two separate sgRNAs or wild-type (WT) controls were analyzed for TREX1 and actin expression by western blotting (**a**) or *ISG15* expression by RT-qPCR (**b**). Data show the relative expression of *IFIT2* in KO cells compared parental controls (set to 1) from n=3 (OE21) or n=6 (SCC25, H596, Detroit 562) replicates, error bars show the SEM. (**c**) Relative levels of cytosolic mtDNA was analyzed following subcellular fractionation by qPCR as detailed in Methods and normalized relative to Calu-3 cells (set to 1). Data show the mean from n=2 replicates, error bars show the SEM. (**d**) SCC25, H596, OE21 and Detroit 562 cells grown in the presence of 100 μM AZT for 3 days. *IFIT1*, *IFIT2* and *IRF7* expression was analyzed by RT-qPCR and normalized relative to mock-treated controls (set to 1). Data show the mean from n=3 replicates, error bars show the SEM. (**e**) RNA expression of the indicated HERV and LINE elements were analyzed in total cell lysates by RT-qPCR (n=2, error bars show the SEM).

**Figure S9. Analysis of ISG and STING expression in SARS-CoV-2-infected primary human airway epithelial cells (hTECs) grown at ALI. (a-c)** hTECs from three donors (Donor 3-5) grown at ALI were mock-infected or infected with SARS-CoV-2-mNG at a MOI of 2 i.u./cell and fixed at 72 hpi. Immunofluorescence detection for ISG15 expression (in red, **a**), STING (in red, **b, c**) mNG (in green), and cellular nuclei (DAPI, in blue) is shown. Images were analyzed by confocal microscopy with a ×40/1.4 objective. Scale bars = 20 and 50μm, (representative of n=2). Mander's overlap coefficient derived from imaging experiments for STING (**d**). Colocalization analysis of Z stacks was performed using the Colocalization module of Volocity from 5-8 separate Z-stacks. Individual numbers plotted display the corresponding values from each Z-stack for each Donor, error bars show the SEM. (**e**) RT-qPCR analysis of ISG expression in hTECs treated with 20 μM SN-011 and infected with SARS-CoV-2 as in Fig.9.

### SUPPLEMENTARY DATA

#### Table S1 sgRNA sequences

| | Target | sgRNA | Reference |
|---|---|---|---|
| 1 | cGAS-gRNA1 (cGAS-1) | ATCCCTCCGTACAAAAAGGG | This paper |
| 2 | cGAS-gRNA2 (cGAS-2) | AGACTCGGTGGGATCCATCG | This paper |
| 3 | STING-gRNA1 (STING-1) | CATATTACATCGGATATCTG | This paper |
| 4 | STING-gRNA2 (STING-2) | CATTACAACAACCTGCTACG | This paper |
| 5 | TBK1-gRNA1 (TBK1-1) | ACAGTGTATAAACTCCCACA | This paper |
| 6 | TBK1-gRNA2 (TBK1-2) | AAATATCATGCGTGTTATAG | This paper |
| 7 | IRF3-gRNA1 (IRF3-1) | CCACTGGTGCATATGTTCCC | This paper |
| 8 | IRF3-gRNA2 (IRF3-2) | AGAAGGGTTGCGTTTAGCAG | This paper |
| 9 | IFNAR-gRNA1 (IFNAR-1) | GTACATTGTATAAAGACCAC | This paper |
| 10 | IFNAR-gRNA2 (IFNAR-2) | ACATGACCTTTCAAGTTCAG | This paper |
| 11 | STAT1-gRNA1 (STAT1-1) | TGCTGGCACCAGAACGAATG | This paper |
| 12 | STAT1-gRNA2 (STAT1-2) | ACAGCAGAGCGCCTGTATTG | This paper |

| | | | |
|---|---|---|---|
| 13 | TREX1-gRNA1 (TREX1-1) | GTCCCCTCCAGACTCGCACA | This paper |
| 14 | TREX1-gRNA2 (TREX1-2) | TCTGGATGGTGCCTTCTGTG | This paper |

**Table S2 Primer sequences**

| **Oligo Target** | **Oligo Sequence** |
|---|---|
| SARS-CoV-2 NP-F | 5’ ATGCTGCAATCGTGCTACAA 3’ |
| SARS-CoV-2 NP-R | 5’ GACTGCCGCCTCTGCTC 3’ |
| IFIT1 (ISG56)-F | 5’ CCTCCCTGGAAAATCTAGGCTCT 3’ |
| IFIT1 (ISG56)-R | 5’ GTAAAGTGACATCTCAATTGCTCCAGAC 3’ |
| IFIT2 (ISG54)-F | 5’ AGCTGAGAATTGCACTGCAACCATG 3’ |
| IFIT2 (ISG54)-R | 5’ CTCCATCAAGTTCCAGGTGAAATGGC 3’ |
| IFIT3-F | 5’ GGGCAGTCATGAGTGAGGTC 3’ |
| IFIT3-R | 5’ AAGTTCCAGGTGAAATGGCA 3’ |
| ISG15-F | 5’ CACAGCCCACAGCCCACAG 3’ |
| ISG15-R | 5’ GCTCAGGGACACCTGGAATTCG 3’ |
| MX1-F | 5’ CACTGCGAGGAGATCGGTTC 3’ |
| MX1-R | 5’ CTGTTCTCCTGCACCTCCTTG 3’ |

| | |
|---|---|
| OAS3-F | 5’ GAAGGAGTTCGTAGAGAAGGCG 3’ |
| OAS3-R | 5’ CCCTTGACAGTTTTCAGCACC 3’ |
| OASL-F | 5’ ACAGATGGGACATCGTTGCT 3’ |
| OASL-R | 5’ TAACCCCTCTGCTCCACTGT 3’ |
| CXCL1-F | 5’ CCCAAACCGAAGTCATAGCCA 3’ |
| CXCL1-R | 5’ TTCTTAACTATGGGGGATGCAG 3’ |
| CXCL2-F | 5’ CGAAAAGATGCTGAAAAATGGC 3’ |
| CXCL2-R | 5’ ACATTAGGCGCAATCCAGGT 3’ |
| CXCL8-F | 5’ TGTGTGTAAACATGACTTCCAAGC 3’ |
| CXCL8-R | 5’ CTTGGCAAAACTGCACCTTCA 3’ |
| CXCL10-F | 5’ AAGTGGCATTCAAGGAGTACCT 3’ |
| CXCL10-R | 5’ ACACGTGGACAAAATTGGCT 3’ |
| CCL20-F | 5’ ATGTCAGTGCTGCTACTCCAC 3’ |
| CCL20-R | 5’ AGCATTGATGTCACAGCCTT 3’ |
| IL1B-F | 5’ GAGCTCGCCAGTGAAATGATG 3’ |
| IL1B-R | 5’ GTGGTCGGAGATTCGTAGCTG 3’ |
| IL6-F | 5’ CCAGAGCTGTGCAGATGAGT 3’ |

| | |
|---|---|
| IL6-R | 5’ GCATTTGTGGTTGGGTCAGG 3’ |
| IL18-F | 5’ CCTCAGACCTTCCAGATCGC 3’ |
| IL18-R | 5’ CTTCTACTGGTTCAGCAGCCA 3’ |
| IRF7-F | 5’ GTCCTGGTGAAGCTGGAACC 3’ |
| IRF7-R | 5’ CAGGGAAGACACACCCTCAC 3’ |
| IRF9-F | 5’ GCCCTACAAGGTGTATCAGTTG 3’ |
| IRF9-R | 5’ TGCTGTCGCTTTGATGGTACT 3’ |
| STAT1-F | 5’ GCTCGTTTGTGGTGGAAAGAC 3’ |
| STAT1-R | 5’ TCTCTCATTCACATCTCTCAACTT 3’ |
| STAT2-F | 5’ CCAGCTTTACTCGCACAGC 3’ |
| STAT2-R | 5’ AGCCTTGGAATCATCACTCCC 3’ |
| IFN-β-F | 5’ AAACTCATGAGCAGTCTGCA 3’ |
| IFN-β-R | 5’ AGGAGATCTTCAGTTTCGGAGG 3’ |
| 18S rRNA-F | 5’ CCGCAGCTAGGAATAATGGA 3’ |
| 18S rRNA-R | 5’ CGGTCCAAGAATTTCACCTC 3’ |
| OAS1-F | 5' GCTCCTACCCTGTGTGTGTGT 3' |
| OAS1-R | 5' TGGTGAGAG GACTGAGGAAGA 3' |

| | |
|---|---|
| BST2-F | 5' CACACTGTGATGGCCCTAAT 3' |
| BST2-R | 5' TGTAGTGATCTCTCCCTC AAGC 3' |
| ELF1-F | 5' GGAGTGCGTCTACTTCTGCC 3' |
| ELF1-R | 5' CAGGCCGTAAGGAGCTGTCT 3' |
| B4GALT5-F | 5' AACAGTITCGGAAAATCAATGGC 3' |
| B4GALT5-R | 5' TTCCTCAGCAGGGCATACCTT 3' |
| ERLIN1-F | 5′ GGGGTTGGTGGCTGTCCTGC 3′ |
| ERLIN1-R | 5′ TAGCCTGGTCCACTGGGGCT 3′ |
| ISG15-F (siRNA expt) | 5' CATCTTTGCCAGTACAGGAGC 3' |
| ISG15-R (siRNA expt) | 5' GGGACACCTGGAATTCGTTG 3' |
| ISG20-F | 5’ CTTCCAGGCACTGAAAGAGG 3’ |
| ISG20-R | 5’ AAGCCGAAAGCCTCTAGTCC 3’ |
| RIG-I-F | 5’ TGTCCACCTTCAGAAGTGTCT 3’ |
| RIG-I-R | 5’ AGCAGGCAAAGCAAGCTCTA 3’ |
| MDA5-F | 5’ AGATGCAACCAGAGAAGATCCA 3’ |
| MDA5-R | 5’ TGGCCCATTGTTCATAGGGT 3’ |
| LGP2-F | 5’TGGGCAAGGCGCAGTTT 3’ |

| | |
|---|---|
| LGP2-R | 5’ ACCGAAGCTCCATTCTGCTC 3’ |
| HERV-T Pol-F | 5’ CCCCTACCCTTTTTGGGG 3’ |
| HERV-T Pol-R | 5’ GTACCCCAGGTAGGAAACTCTGGG 3’ |
| HERV-E Pol-F | 5’ CTTTGGGGAGGCGTTGGCTCGAGACC 3’ |
| HERV-E Pol-R | 5’ GCTTTCTTTCTGATCCTAGGCTGTG 3’ |
| ERV9 Pol-F | 5’ CCCTCATCTGTTTGGTCAGGCCC 3’ |
| ERV9 Pol-R | 5’ CCTCAACTGTTTTAATGTCTTAGGGCGAGG 3’ |
| ERV9-Seq59 Pol-F | 5’ CAGGCACAGGCCCAAGATCTAGTTC 3’ |
| ERV9-Seq59 Pol-R | 5’ GTGCTGAGGGCCCTGGTTCCTCTGG 3’ |
| HERV-K (HML-10) Pol-F | 5’ CCCACAGTTTGTCAAACTTTTGTAGGC 3’ |
| HERV-K (HML-10) Pol-R | 5’ GAATCTCTTCTAATTTGAACCTTTTGAGG 3’ |
| HERV-K (HML-2) Gag-F | 5’ GGCCATCAGAGTCTAAACCACG 3’ |
| HERV-K (HML-2) Gag-R | 5’ CTGACTTTCTGGGGGTGGCCG 3’ |
| SINE B1-F | 5’ TGGCGCACGCCTTTAATC 3’ |
| SINE B1-R | 5’ TGGCCTCGAACTCAGAATCC 3’ |
| LINE1 5’ UTR-F | 5’ TAAACAAAGCGGCCGGGAA 3’ |
| LINE1 5’UTR-R | 5’ AGAGGTGGAGCCTACAGAGG 3’ |

| | |
|---|---|
| LINE1 ORF1-F | 5’ ACCTGAAAGTGACGGGGAGA 3’ |
| LINE1 ORF1-R | 5’ CCTGCCTTGCTAGATTGGGG 3’ |
| LINE1 ORF2-F | 5’ CAAACACCGCATATTCTCACTCA 3’ |
| LINE1 ORF2-R | 5’ CTTCCTGTGTCCATGTGATCTCA 3’ |
| KCNJ-F | 5’ GCGCAAAAGCCTCCTCATT 3’ |
| KCNJ-R | 5’ CCTTCCTTGGTTTGGTGGG 3’ |
| MT-ND1-F | 5’ GAACTAGTCTCAGGCTTCAACATCG 3’ |
| MT-ND1-R | 5’ CTAGGAAGATTGTAGTGGTGAGGGTG 3’ |
| MT-D-loop-F | 5’ CATAAAGCCTAAATAGCCCACACG 3’ |
| MT-D-loop-R | 5’ CCGTGAGTGGTTAATAGGGTGATA 3’ |

**Table S3 siRNA information from Ambion by Life Technologies**

| Target | Sequence | Identifier |
|---|---|---|
| siTLR3 - 1 | Sense: CCGAUGAUCUACCCACAAAtt<br>Antisense: UUUGUGGGUAGAUCAUCGGgt | s236 |
| siTLR3 – 2 | Sense: GGAUAGGUGCCUUUCGUCAtt<br>Antisense: UGACGAAAGGCACCUAUCCgt | s235 |
| siTLR7 - 1 | Sense: GGAAUUACUCAUAUGCUAAtt<br>Antisense: UUAGCAUAUGAGUAAUUCCtt | s27842 |
| siTLR7 - 2 | Sense: CCAUUAACCACAUACCAGAtt<br>Antisense: UCUGGUAUGUGGUUAAUGGtg | s27843 |
| siTLR8 - 1 | Sense: GGAUCUUGAAUUCAACUAUtt<br>Antisense: AUAGUUGAAUUCAAGAUCCag | s27921 |
| siTLR8 - 2 | Sense: GACCCAACUUCGAUACCUAtt<br>Antisense: UAGGUAUCGAAGUUGGGUCaa | s27920 |
| siTLR9 - 1 | Sense: ACAAUAAGCUGGACCUCUAtt<br>Antisense: UAGAGGUCCAGCUUAUUGUgg | s28873 |
| siTLR9 - 2 | Sense: CUGGAAGAGCUAAACCUGAtt<br>Antisense: UCAGGUUUAGCUCUUCCAGgg | s28872 |
| siRIG-I - 1 | Sense: GGAUUGUUACAGUUCAGAAtt<br>Antisense: UUCUGAACUGUAACAAUCCat | s223616 |
| siRIG-I - 2 | Sense: GAAGCAGUAUUUAGGGAAAtt<br>Antisense: UUUCCCUAAAUACUGCUUCgt | s223615 |
| siMDA5 - 1 | Sense: GGUGUAAGAGAGCUACUAAtt<br>Antisense: UUAGUAGCUCUCUUACACCtg | s34499 |
| siMDA5 - 2 | Sense: GUUCAGGAGUUAUCGAACAtt<br>Antisense: UGUUCGAUAACUCCUGAACca | s34500 |
| siLPG2 - 1 | Sense: GGCAGUUCAGUAGCUCUAAtt<br>Antisense: UUAGAGCUACUGAACUGCCtt | s35594 |
| siLPG2 - 2 | Sense: CCGGAAAUUUGGGACGCAAtt<br>Antisense: UUGCGUCCCAAAUUUCCGGct | s35595 |
| siMyD88 - 1 | Sense: CGGCAACUGGAACAGACAAtt<br>Antisense: UUGUCUGUUCCAGUUGCCGga | s9137 |

| | | |
|---|---|---|
| siMyD88 - 2 | Sense: CAGACAAACUAUCGACUGAtt<br>Antisense: UCAGUCGAUAGUUUGUCUGtt | s9138 |
| siTRIF – 1 | Sense: GAAUCAUCAUCGGAACAGAtt<br>Antisense: UCUGUUCCGAUGAUGAUUCca | s45114 |
| siTRIF - 2 | Sense: CAACUUUCUGUAGAAGAUAtt<br>Antisense: UAUCUUCUACAGAAAGUUGga | s531858 |
| siMAVS - 1 | Sense: CUGUUGCAUUGGUCCCUGAtt<br>Antisense: UCAGGGACCAAUGCAACAGga | s33179 |
| siMAVS – 2 | Sense: GGGUUCUUCUGAGAUUGAAtt<br>Antisense: UUCAAUCUCAGAAGAACCCag | s33180 |
| siNOD2 - 1 | Sense: GCACAACCUUCAGAUCACAtt<br>Antisense: UGUGAUCUGAAGGUUGUGCgg | s533471 |
| siNOD2 - 2 | Sense: GGAACACUUUCUCUCUAGAtt<br>Antisense: UCUAGAGAGAAAGUGUUCCct | s34485 |
| siNLRP1 - 1 | Sense: GUACGAGACUCGGAACAAAtt<br>Antisense: UUUGUUCCGAGUCUCGUACaa | s22520 |
| siNLRP1 - 2 | Sense: GGUGGAGCUGCAUCACAUAtt<br>Antisense: UAUGUGAUGCAGCUCCACCct | s22521 |
| siNLRP1 - 3 | Sense: CGGUGACCGUUGAGAUUGAtt<br>Antisense: UCAAUCUCAACGGUCACCGct | s22522 |
| siNLRP3 - 1 | Sense: GAGAGACCUUUAUGAGAAAtt<br>Antisense: UUUCUCAUAAAGGUCUCUCct | s534395 |
| siNLRP3 - 2 | Sense: GCACGUGUUUCGAAUCCCAtt<br>Antisense: UGGGAUUCGAAACACGUGCat | s41556 |
| siNLRC5 - 1 | Sense: GCUGAUCUUUGAUGGGCUAtt<br>Antisense: UAGCCCAUCAAAGAUCAGCag | s38591 |
| siNLRC5 - 2 | Sense: GAGCUGGACUUGUCUAACAtt<br>Antisense: UGUUAGACAAGUCCAGCUCct | s38592 |
| siSTING - 1 | Sense: GGUCAUAUUACAUCGGAUAtt<br>Antisense: UAUCCGAUGUAAUAUGACCat | s50644 |
| siSTING - 2 | Sense: CCUCAUCAGUGGAAUGGAAtt<br>Antisense: UUCCAUUCCACUGAUGAGGag | s226307 |
| siSTAT2 - 1 | Sense: GGCUCAUUGUGGUCUCUAAtt<br>Antisense: UUAGAGACCACAAUGAGCCtg | s13528 |

| | | |
|---|---|---|
| siSTAT2 - 2 | Sense: CAACAACCGCUGGAGCUUAtt<br>Antisense: UAAGCUCCAGCGGUUGUUGca | s13529 |
| siSTAT2 - 3 | Sense: GGCCGAUUAACUACCCUAAtt<br>Antisense: UUAGGGUAGUUAAUCGGCCta | s13530 |
| siIRF9 - 1 | Sense: CACAGAAUCUUAUCACAGUtt<br>Antisense: ACUGUGAUAAGAUUCUGUGga | s20290 |
| siIRF9 - 2 | Sense: CAAUAUUUAAGGGAAAGUAtt<br>Antisense: UACUUUCCCUUAAAUAUUGcc | s20291 |
| siIRF9 - 3 | Sense: GCUAAGACCAUGUUCCGGAtt<br>Antisense: UCCGGAACAUGGUCUUAGCtg | s20292 |
| ELF1-1 | Sense: GGUCCUUACUAGCAAUGUUtt<br>Antisense: AACAUUGCUAGUAAGGACCtg | s4618 |
| ELF1-2 | Sense: GGACUGUUCAUGUAGUACAtt<br>Antisense: UGUACUACAUGAACAGUCCgg | s4617 |
| B4GALT5-1 | Sense: GGGACAACGUGAGAACAAUtt<br>Antisense: AUUGUUCUCACGUUGUCCCgg | s17844 |
| B4GALT5-2 | Sense: CAACCAAAUUGGAUAAGUAtt<br>Antisense: UACUUAUCCAAUUUGGUUGca | s17845 |
| ERLIN1-1 | Sense: GAUUAUGACAAGACCUUAAtt<br>Antisense: UUAAGGUCUUGUCAUAAUCtg | s20840 |
| ERLIN1-2 | Sense: GGAAUAUCUGGAGCUCAAAtt<br>Antisense: UUUGAGCUCCAGAUAUUCCgg | s20839 |
| ISG15-1 | Sense: CCAUGUCGGUGUCAGAGCUtt<br>Antisense: AGCUCUGACACCGACAUGGag | s18524 |
| ISG15-2 | Sense: GCACCGUGUUCAUGAAUCUtt<br>Antisense: AGAUUCAUGAACACGGUGCtc | s195001 |
| BST2-1 | Sense: AUCACUACAUUAAACCAUAtt<br>Antisense: UAUGGUUUAAUGUAGUGAUct | s2104 |
| BST2-2 | Sense: GGAAUUCUGGUGCUCCUGAtt<br>Antisense: UCAGGAGCACCAGAAUUCCta | s2103 |
| OAS1-1 | Sense: CCACUUUUCAGGAUCAGUUtt<br>Antisense: AACUGAUCCUGAAAAGUGGtg | s511 |
| OAS1-2 | Sense: GUCAAGCACUGGUACCAAAtt<br>Antisense: UUUGGUACCAGUGCUUGACta | s224141 |

| | | |
|---|---|---|
| ISG20-1 | Sense: AGAGAUCACCGAUUACAGAtt<br>Antisense: UCUGUAAUCGGUGAUCUCUcc | s7524 |
| ISG20-2 | Sense: AGCUGGUGGUGGGUCAUGAtt<br>Antisense: UCAUGACCCACCACCAGCUtg | s7523 |

**Figure S1B-Uncropped western blot images:**

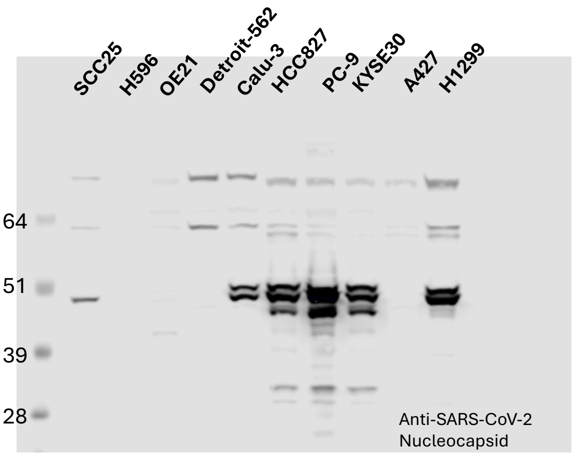


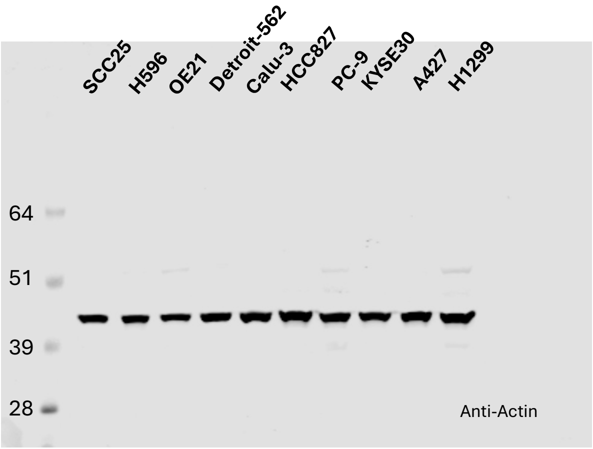


**Figure S3D-Uncropped western blot images:**

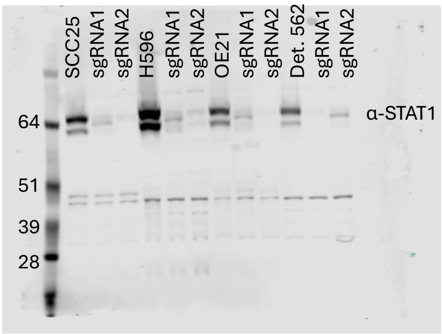


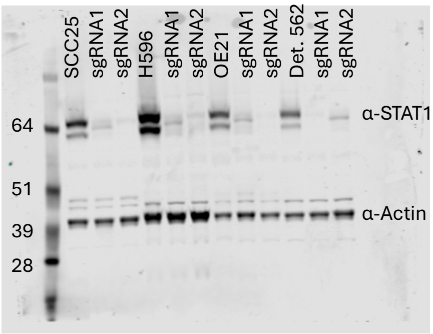


**Figure S4B-Uncropped western blot images:**

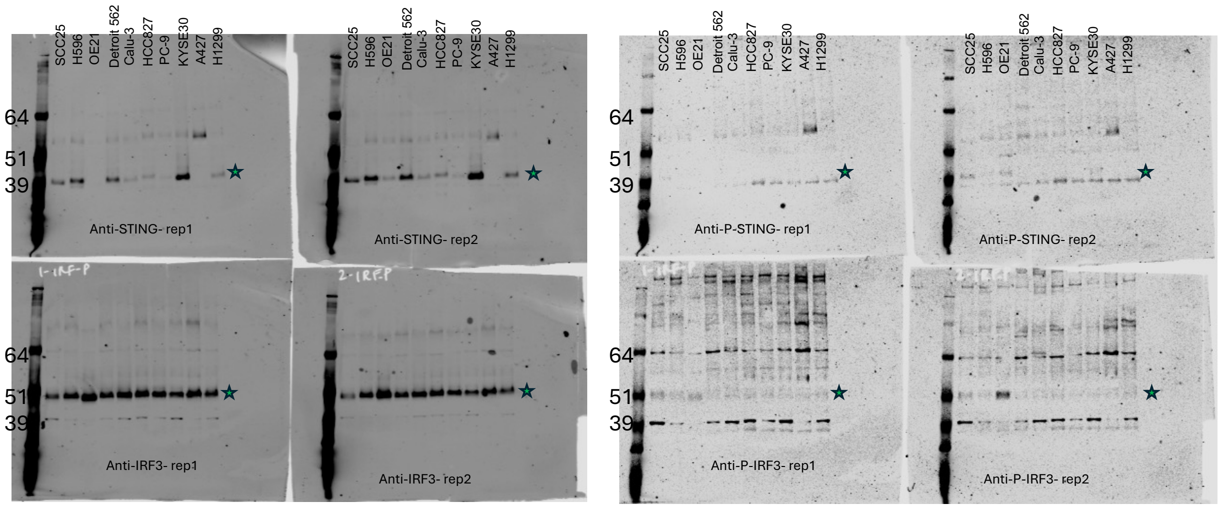


### Figure S4C-Uncropped western blot images:

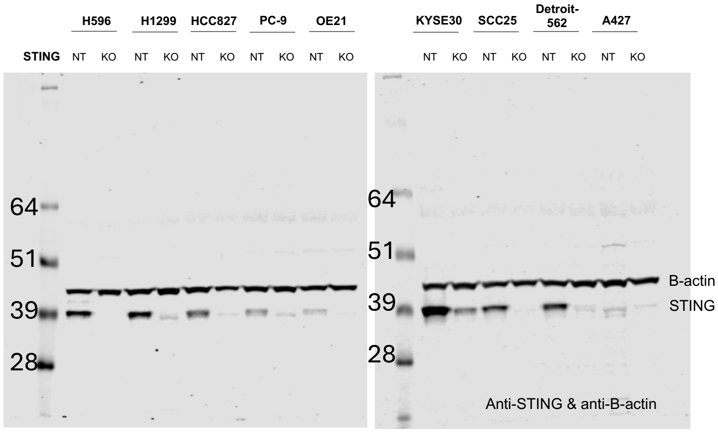


### Figure S8A-Uncropped western blot image:

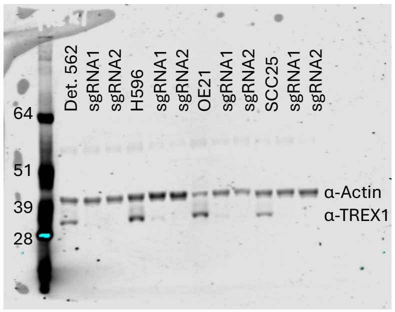
